# A single-cell atlas of the human brain in Alzheimer’s disease and its implications for personalized drug repositioning

**DOI:** 10.1101/2022.06.14.496100

**Authors:** Guangsheng Pei, Brisa S Fernandes, Yin-Ying Wang, Astrid M Manuel, Peilin Jia, Zhongming Zhao

**Affiliations:** Center for Precision Health, School of Biomedical Informatics, The University of Texas Health Science Center at Houston, Houston, TX 77030, USA; Human Genetics Center, School of Public Health, The University of Texas Health Science Center at Houston, Houston, TX 77030, USA; Department of Biomedical Informatics, Vanderbilt University Medical Center, Nashville, TN 37203, USA

**Keywords:** Alzheimer’s disease, subpopulation, cell-cell communication, ligand-receptor, drug repositioning

## Abstract

Alzheimer’s disease (AD) is a neurodegenerative disease with complex pathophysiology, and AD-dysregulated pathways are inconsistent across different brain regions and patients. Although single-cell RNA sequencing (scRNA-seq) has been performed in different regions of postmortem AD brains, the common and distinct molecular features among different regions remains largely unclear. This hinders the discovery of repurposable and personalized drugs for AD. We combined four scRNA-seq datasets and systematically investigated the common and distinct cellular responses, cell subpopulations, and transcription factors involved in AD. Moreover, we explored the transcriptional heterogeneity of different AD subtypes at the single-cell level. Finally, we conducted individual-based drug repurposing analysis to explore repurposable and personalized drugs. Six major brain cell types were detected after scRNA-seq batch-effect removal and noise cells filtering. Integration with genome-wide association studies (GWAS) summary statistics demonstrated that AD-susceptible genes were mainly enriched with differentially expressed genes (DEGs) in glial cells rather than neuronal cells. While most of DEGs were regulated in opposite directions among different cell types, cell-cell communication analysis revealed several common cellular interaction events involved in neurogenesis, as well as increased cell-cell adhesion. Our comprehensive drug repositioning analysis identified new candidates for AD treatment, including trichostatin, which was predicted to be broadly applicable to different identified AD subtypes, and vorinostat, which was specific for one subtype of AD. In summary, we delineated a cell-specific atlas of the AD transcriptome. Our work illustrated strong cellular heterogeneity in AD for defining AD subtypes. The cell-specific features are important for understanding AD etiology, progression, and drug discovery.

## Introduction

Alzheimer’s disease (AD) is the most common form of dementia and has significant individual and socio-economic burdens [1, 2]. The pathogenesis of AD is heterogeneous and complex, with a variety of risk factors contributing to its occurrence, such as family history, lifestyle, environment, and aging [2]. Two decades ago, researchers classified AD into two main types. The so-called late-onset AD (LOAD) is the predominant type of AD (approximately 95% of all cases), which usually occurs after the age of 65, is sporadic without an obvious inheritance pattern. Another type of AD, which accounts for less than 5% of all AD cases, is a rare familial form of AD and appears to be genetically inherited, typically occurring earlier than age 65, named early-onset AD (EOAD) [2, 3]. Our current understanding is that AD, regardless if it is early or late-onset, is primarily driven by the extracellular deposition of amyloid-beta (Aβ) peptide as extracellular plaques. AD is also characterized by the presence of hyper-phosphorylated tau protein as intracellular neurofibrillary tangles (NFTs) and consequent neurodegeneration, in tandem with downstream inflammatory processes [2, 4]. Despite this singular proposed pathophysiology, the progression and treatment response in LOAD is highly heterogeneous; this challenges our contemporary conceptualization of AD as a disease with unitary biology and suggests the presence of distinct pathophysiologic subtypes [4]. The presence of subtypes could also explain why medications prescribed to delay the progression of AD symptoms are not efficacious in all individuals with AD [2].

Characterizing the cellular heterogeneity and transcriptional changes in the brain microenvironment of AD people may likely help to disentangle the complex pathophysiology of AD. To advance on this front, single cell and single-nucleus RNA-sequencing (scRNA-seq and snRNA-seq) studies have been done in different regions of postmortem brains of individuals with AD. However, to fully leverage these laboratory techniques and to systematically chart the landscape of cellular diversity in AD, robust bioinformatics methods are necessary. Previous works studied the cell-type spatiotemporal specificity through human brain development [5]. However, most of those studies employed bulk RNA; in those studies, different brain regions or developmental stages were found to have distinct vulnerabilities to AD [6]. To make sense of differences in cell-type spatiotemporal specificity throughout human brain development, the delineation of a bioinformatics-informed atlas of multi-brain regions in AD using single cell data is necessary. This approach will also facilitate the understanding of the cell population’s regional specificity and selective vulnerability. Secondly, the development and progression of AD involve finely tuned cascades of interactive events. For instance, neural cells not only communicate to impart particular cell fates upon each other [7], but also communicate with developing vascular cells and microglia to guide their development [8]; microglia, the resident immune cells of the central nervous system, are observed around NFTs in AD brains [9]. Despite accumulating evidence highlighting interactions between neural cells and glial cells as critically important for neurogenesis in AD [7, 10], the extent of dysregulated communication patterns among major brain cell types in AD remains to be addressed. Finally, recent studies highlighting molecular features in bulk RNA-seq at the group-level even have opposite directions of differential gene expression in different AD subtypes, which may be due to AD’s hypothesized heterogeneity [4]. Therefore, single-cell level investigations might offer more efficacious personalized AD treatments.

In this study, we conducted an integrative analysis of four existing scRNA-seq and snRNA-seq datasets, as well as two bulk RNA-seq cohorts across a spectrum of AD brains. After assembling existing scRNA-seq/snRNA-seq datasets from different resources, we characterized the cell subpopulation diversity between individuals with AD and healthy controls. Leveraging dysregulated communication patterns of AD among different brain regions and severity, we further shaped the landscape in AD. We subsequently demonstrated substantial heterogeneity among existing AD subtypes at both single-cell and bulk levels, indicating distinct pathogenic mechanisms and advancing the knowledge of AD heterogeneity, which may be applicable for precision medicine. Finally, individual-level drug repurposing analysis allowed us to identify histone deacetylase inhibitors – trichostatin and vorinostat –, as candidates for drug repurposing in AD.

## Methods

### AD transcriptome cohorts

#### Single-cell RNA-seq data

AD scRNA-seq profiles were downloaded from scREAD (single-cell RNA-Seq database for AD [11]) and Synapse (syn18485175, February 25, 2021). In total, there were more than 300,000 cells from four different human AD scRNA-Seq/snRNA-Seq data repositories [12–15], covering three different brain regions [prefrontal cortex (PFC, Brodmann areas 9, 10, and 46), entorhinal cortex (EC), and superior frontal gyrus (SFG, Brodmann area 8)]. A total of eight different cell types were identified (astrocytes, excitatory and inhibitory neurons, microglia, oligodendrocytes, oligodendrocyte precursor cells (OPCs), endothelial cells, and pericytes). Due to very small proportions, endothelial cells and pericytes were excluded in the downstream analysis. See Additional file 1: Tables S1 and S2 for more details on scRNA-seq/snRNA-Seq data. In addition, the clinical information of snRNA-seq datasets in prefrontal cortex region, *e.g.*, level of AD pathology burden (PB) and amyloid level, was collected [13].

### Bulk RNA-seq data

The AD bulk RNA-seq data was downloaded from the Mount Sinai Brain Bank (MSBB) [16] and the Religious Orders Study/Memory and Aging Project (ROSMAP) [17, 18] databases (April 29, 2021). The MSBB-AD cohort included 938 RNA expression data stratified by clinical dementia rating (CDR) score [control: CDR = 0; mild cognitive impairment (MCI): CDR = 0.5; AD brain: CDR ≥ 1] and four different brain regions, including: frontal pole cortex (FP, Brodmann area 10, *N_Ctr_* = 35, *N_MCI_* = 39, *N_AD_*=187), inferior frontal gyrus (IFG, Brodmann area 44, *N_Ctr_* = 27, *N_MCI_* = 37, *N_AD_ =* 157), parahippocampal gyrus (PHG, Brodmann area 36, *N_Ctr_* = 32, *N_MCI_* = 32, *N_AD_* =151), and superior temporal gyrus (STG, Brodmann area 22, *N_Ctr_*= 33, *N_MCI_* = 33 *N_AD_*= 174). The ROSMAP cohort [17, 18] included 638 transcriptome stratified by cognitive diagnosis score from the dorsolateral prefrontal cortex (DLPFC). After excluding the samples unrelated to AD, a total of 257 brains of deceased AD individuals, 168 brains of deceased MCI individuals, and 201 brains of healthy controls were included in the downstream analyses.

### scRNA-seq/snRNA-seq data integration

After obtaining the raw count matrix from scREAD [11] and Synapse [13], the R package Seurat (v4.0.3) [19] was used for downstream analysis on the R platform environment (v4.0.2). To align different data batches, we utilized Seurat reciprocal principal component analysis (PCA) based integration (function: *FindIntegrationAnchors*) to determine the anchor genes between different datasets. Compared to canonical correlation analysis [20], reciprocal PCA employs a more conservative approach to avoid overcorrection, and it also runs significantly faster. In addition, batch-specific cell types (pericytes and endothelial cells) and transcription noise cells were also filtered by several criteria, including minimal expression of 300 genes per cell and mitochondrial read percentage >30%. The cell cycle phase analysis was then scored based on its expression of phase markers [21]. All cells passing quality control were merged into one count matrix, normalized and scaled using Seurat’s *NormalizeData* and *ScaleData* functions. The reduced set of highly variable genes was used as the feature set for independent component analysis on 3000 genes using Seurat’s RunPCA function. A Uniform Manifold Approximation and Projection (UMAP) dimensional reduction analysis [22] was performed on the scaled matrix (with only the most variable genes) using the first 30 PCA components to obtain a two-dimensional representation of the cell states. Then, cell clustering was conducted using the function *FindClusters,* which implements the shared nearest neighbor modularity optimization-based clustering algorithm with resolution = 0.8, leading to 33 cell clusters. The differentially expressed gene (DEG) analysis between clusters was performed using the Wilcoxon rank-sum test. After detecting the top hits for each cluster, cell types were annotated manually based on prior knowledge and the *deCS* tool [23].

### DrivAER analysis to identify core transcription factors

Due to the complexity and multifactorial AD pathophysiological characterization, initial efforts using a conventional fold change has led to the determination of DEGs at the single cell level [12–15]. However, it became clear that data manifold methods, rather than individual gene-focused methodologies, would be more appropriate to investigate subtle changes of transcriptional signals on single cells. Here, we applied **Driv**ing transcriptional programs based on **A**uto**E**ncoder derived **R**elevance scores (DrivAER) [24], which identifies driving transcriptional programs in single-cell RNA sequencing data. It employs a Deep Count Auto-encoder and random forest (RF) model to identify transcriptional programs in AD pathology. We downloaded the transcriptional program annotations from The Molecular Signatures Database (MSigDB) [23] and the Transcriptional Regulatory Relationships Unraveled by Sentence-based Text-mining (TRRUST) [24]. For each cell type, the top 10 transcription factors (TF) between the brains of AD individuals and healthy controls were identified.

### Cell-cell communication analysis

AD may affect brain cellular interactions through an interactive connection among cell types, including neurons, astrocytes, and microglia [10]. The brain comprises several regions and various cell types [5]. To identify and visualize the cell state-specific cell-cell interactions, we employed an R package called CellChat [25] to infer AD cell-to-cell interactions in distinct neocortical areas or AD stages. Briefly, we loaded the normalized counts into CellChat, and applied the standard preprocessing steps, which involved the application of the functions *identifyOverExpressedGenes*, *identifyOverExpressedInteractions*, and *projectData* with default parameter settings. A total of 2,021 pre-validated ligand-receptor (L-R) interactions were selectively used as a priori network information. For each L-R pair, we then calculated their information flow strength and communication probability between different cell groups by using the functions *computeCommunProb*, *computeCommunProbPathway*, and *aggregateNet* with standard parameters [25]. Together, the overall communication probabilities among all pairs of cell groups across all pairs of L-R interactions were transformed into a three-dimensional tensor *P*, (*K* × *K* × *N*), where *K* corresponds to six cell groups, and *N* corresponds to L-R pairs of different signaling pathways [25].

To predict significant intercellular communications between the control and AD groups, for each L-R pair, we used a one-sided permutation test (n = 100), which randomly permuted the group labels of cells and then recalculated the communication probability between two cell groups [25]. The interactions with a *p*-value <0.05 were considered statistically significant.

### Bulk RNA-seq deconvolution analysis

To evaluate the cell type composition among different molecular subtypes of AD [4] from the MSBB-AD [16] and the ROSMAP [17, 18] cohorts, we applied Scaden (Single-cell assisted deconvolutional network) [26], a deep neural network model, to estimate the relative cellular fraction of six human brain cell types. To balance the number of cells, we randomly selected 100 cells in each cell type as the reference panel.

### AD-associated genes from genome-wide association study (GWAS)

We downloaded AD-associated GWAS datasets from a recent meta-analysis of AD [27]. To define AD susceptible genes, we mapped SNPs to genes if they were located in the gene body or 50 kb upstream and 35 kb downstream of the gene using the Multi-marker Analysis of GenoMic Annotation (MAGMA) software tool [28]. Then, we used the mean Chi-square statistic for these SNPs to obtain the gene-based *p*-value (Details in Additional file 1: Table S3), considering the effects of the gene length, SNP density, and local linkage disequilibrium structure. To ensure the reliability, we stratified AD susceptible genes by using different thresholds: from 10^-4^ to 10^-8^.

### Functional pathway and spatiotemporal analysis of AD differentially expressed genes

The differential expression analysis between different cell clusters was conducted by a Wilcoxon rank-sum test implemented in Seurat ‘*FindAllMarkers’* function. For each cluster, we defined DEGs as those genes that were expressed in more than 25% of cells, had log fold-change greater than 0.25 compared to the background, and the false discovery rate (FDR) were less than 0.05. The distribution of DEGs count was conducted by the UpSetR package (v1.4.0). Functional enrichment analyses of Gene Ontology (GO) and Kyoto Encyclopedia of Genes and Genomes (KEGG) pathways were conducted by the WebGestalt package (v 0.4.4) [29]. All human protein-coding genes were used as the background gene set. Benjamini-Hochberg’s procedure [30] was used for multiple test correction. Significant pathways were defined as those with FDR < 0.01.

We conducted tissue-specific enrichment analysis using the *deTS* method [31–33] on the Genotype-Tissue Expression (GTEx) panel. In addition, the association analysis between spatiotemporal brain development and DEGs of AD was conducted using Fisher’s exact test on 29 gene modules from our previous imputed BrainSpan consortium [5].

### AD associated genes - based classifier

In order to investigate our finding from snRNA-seq on bulk level, 10-fold cross-validation was performed on a random forest model (randomForest package, v4.6-14), based on the transcriptome profiles of the brains of participants with AD, MCI, or healthy controls. The cross-validation error curves from 10 trials of the 10-fold cross-validation were averaged, and the minimum error in the averaged curve plus the standard deviation at that point was used as the cutoff value [34]. All sets of DEG markers with an error less than the cutoff were listed, and the set with the best performance was chosen as the optimal set for downstream prediction. The probability of a brain belonging to a participant with AD or to a healthy control was calculated using this set of conserved DEG markers or the union with dysregulated L-R features. The area under the receiver operating characteristic (AUROC) that was drawn by the pROC package (v1.17.0.1) was used to evaluate the performance. The model was further tested on the test set, and the prediction error was determined on the MSBB-AD and ROSMAP cohorts, respectively.

### Drug repurposing analysis

To predict potentially repurposable drugs for AD, we applied the strategy performed in CeDR [35] to conduct the cellular drug response analysis, which provides references for new therapeutic development and drug combination design at single cell resolution. To this end, we downloaded the drug-induced gene expression matrix from CMap database (version: build 02) which measures 1309 FDA approved drugs with different doses across five cell lines, yielding a total of 6100 profiles [34]. The matrix was ranked based on the DEGs (drug treated versus untreated) and each probe was subsequently mapped to gene symbols. Each scRNA-seq/snRNA-seq dataset was processed followed by the pipeline illustrated in *Scanpy* package [36]. Applying the “anti-correlation” procedure, we defined the cellular gene signatures by combining the top 250 and bottom 250 genes across cell types and within cell type. We further required the drug up-regulated genes to be enriched in cellular down-regulated genes, and vice versa [35]. Moreover, the expression of overlapping genes should be significantly anti-correlated. We independently conducted two Chi-square tests for enrichment analysis and employed the *Spearman* correlation coefficient for anti-correlation. The drugs with significant *p*-values were subsequently denoted as AD cell type-associated drug candidates.

To simplify and keep a highly confident relationship between patients and drug candidates, a patient–drug associated network was constructed after combining the patient–drug association across different cell types. Subsequently, the relationships between different AD individuals were imputed based on the Kendall rank correlation coefficient.

## Results

### Variable cell type composition changes in AD brains and their regions

Our goal is to decode the cellular heterogeneity and microenvironment changes in AD brain and inference for personalized drug repositioning (Figure 1). To this end, we first investigated the cell diversity and cellular changes in AD to illustrate their common or region-specific features. We applied reciprocal PCA [20] to four existing assembled human AD scRNA-seq/snRNA-seq datasets covering three different brain regions (prefrontal cortex, PFC; entorhinal cortex, EC; superior frontal gyrus, SFG). After reprocessing the scaled gene-barcode matrices, low-quality cells removal and standard clustering analysis in Seurat [20], a total of 33 distinct clusters were identified by canonical marker genes (Figure 2A-C). These clusters comprised six major brain cell types, astrocytes (*AQP4*), excitatory (*SLC17A7*), and inhibitory neurons (*GAD2*), microglia (*CD74*), oligodendrocytes (*VCAN*), and OPCs (*MOG*), The UMAP analysis confirmed that the batch effects were minimized among different brain regions. Subdividing the total pool of cells into two categories based on the disease status of the donors (AD brain/healthy control) showed that both conditions appeared in all clusters (Figure 2D, Additional file 2: Figure S1). Then, the proportions of broad cell type changes among different regions were assessed. We observed an overall downward trend in the relative abundance of excitatory neurons in the SFC and EC region (Figure 2E) [14]. However, most cell type fractions in the PFC region did not reach statistical significance [13]. In contrast, we observed that the fraction of astrocytes, oligodendrocytes, and microglia increased with different extents in different brain regions (Figure 2F). To better investigate this finding, we further performed cell type deconvolution analysis from two large bulk cohorts, the MSBB AD [16] and the ROSMAP [17, 18], across five brain regions. We observed decreased patterns of excitatory neurons and increased patterns of astrocytes and oligodendrocytes across all brain regions (Additional file 2: Figure S2). Lastly, we estimated the signature difference of the cell cycle phase among different brain regions (Figure 2G). We observed that the proportion of cells on the S phase decreased, while the proportion on the G1 phase increased (Figure 2H) in the SFC and EC, suggesting a failure of the cells to undergo a new cycle division and DNA synthesis to start transcription [37]. Collectively, our results indicated that AD pathology affect brain regions and cells in an un-uniformly manner [6, 38].

**Figure 1.**
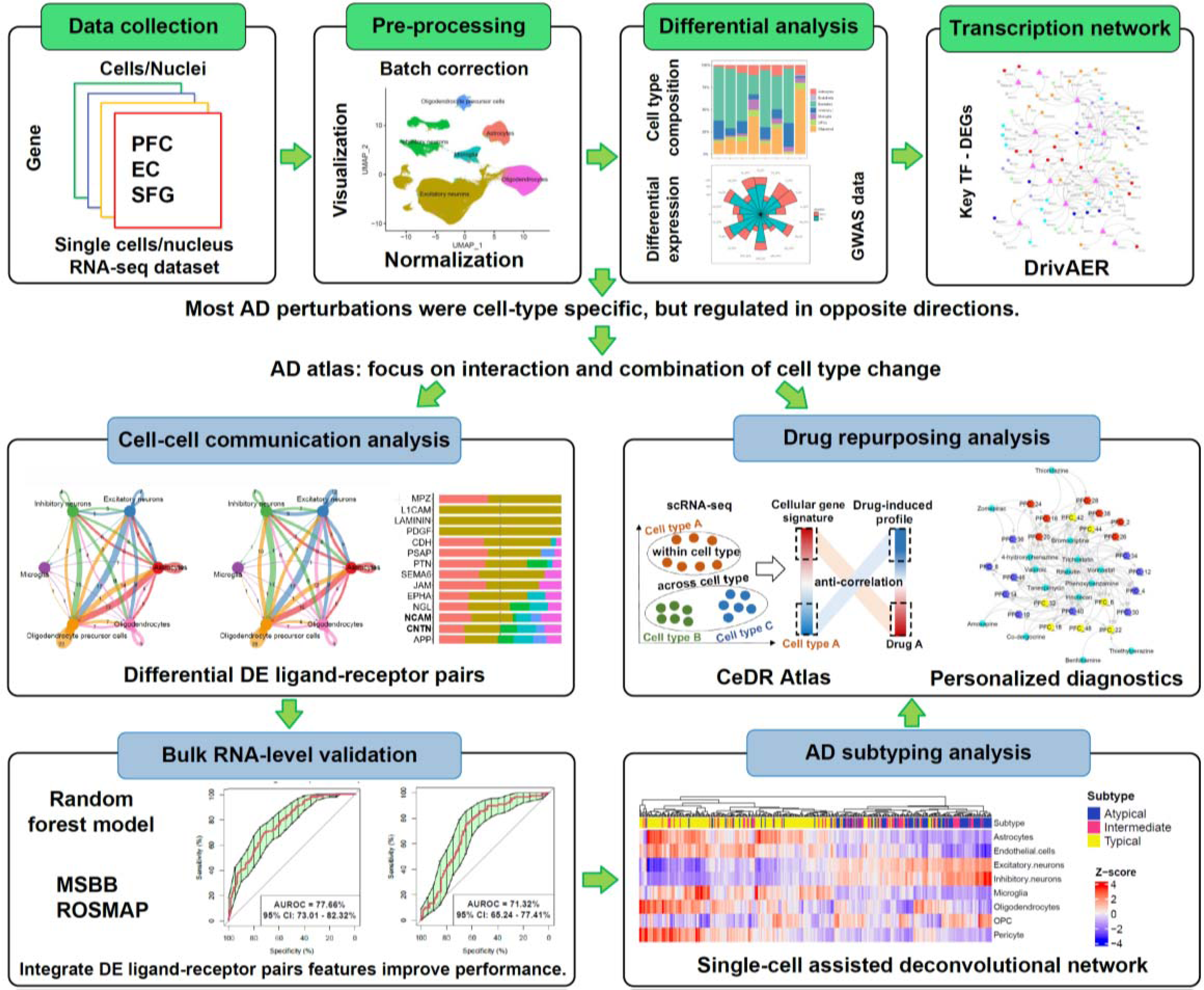
The workflow of the study. Details are provided in the Methods section. We first performed a comprehensive analysis of scRNA-seq/snRNA-seq datasets across three brain regions. Next, we sought to determine if there were any unexpected conserved intercellular communication patterns in AD. Finally, we conducted AD subtyping analysis and personalized drug repurposing analyses based on the single-cell atlas of the human brain in AD.

**Figure 2.**
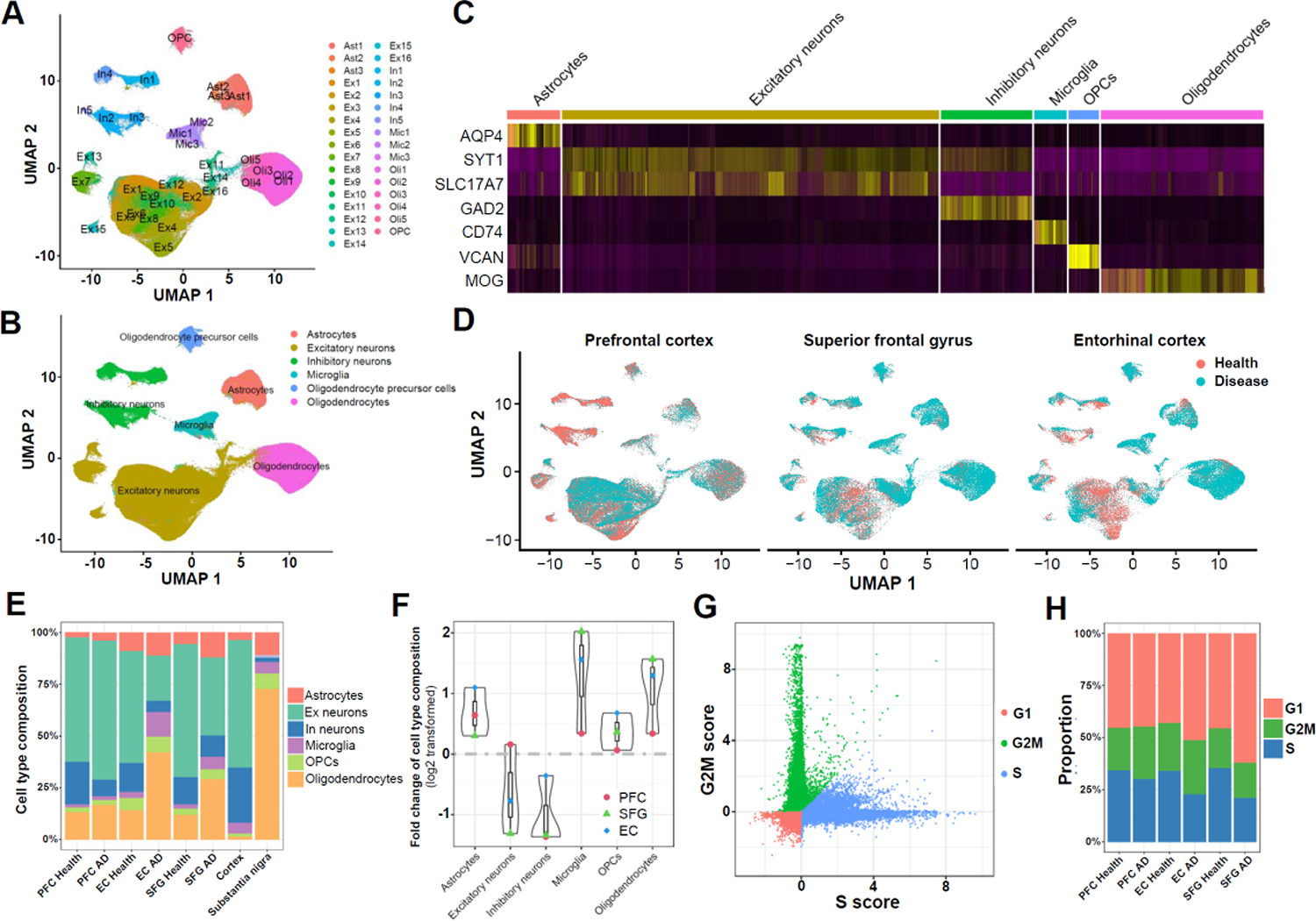
Single-cell RNA-seq integrative analysis among different AD brain regions. **(A)** UMAP projection of 33 cell clusters were annotated into **(B)** six major cell types from the PFC, EC, and SFG. **(C)** Heat map of canonical marker gene expression across different cell types. **(D)** UMAP breakdown according to the cohort of origin regions indicated that our integrative analysis successfully overcame batch effects across different cohorts/platforms. Cells are color-coded by their corresponding health conditions. **(E)** The relative cell-type composition of health and AD brains among PFC, EC, and SFG regions (the cell type composition of cortex and substantia nigra was based on Agarwal *et al*. 2020). **(F)** The progressive pattern of AD pathology among different cell types after log2 transformation. **(G)** The cell cycle score distribution and **(H)** the percentage of cells in G1, G2/M, and S phases among PFC, EC, and SFG.

### The common perturbations at different brain regions were cell-type specific but regulated in divergent direction

Although previous studies demonstrated that the up-and downregulated DEGs in AD brains were highly concordant [12–15], different DEGs were found in different brain regions [14]. Therefore, we conducted a comprehensive comparison of DEGs analysis between AD and healthy control. As shown in Figure 3A, most cell types in the EC region had the largest median number of DEGs (1485 genes, FDR < 0.05) compared to the PFC (844 genes) and the SFG (471 genes). In addition, approximately 60% of the DEGs in the PFC region were mainly enriched in excitatory and inhibitory neurons. Collectively, we obtained a total of 532 DEGs regulated in the same direction in the three brain regions, PFC, EC, and SFG (co-directional), in at least one cell type (defined as conserved DEGs) (Additional file 1: Table S4). Although 54.9% (292/532) DEGs overlapped in multiple cell types, only 25% of DEGs demonstrated co-directional overlap among PFC, EC, and SFG in at least one cell type (Figure 3B). Most conserved DEGs were regulated in opposite directions among different cell types. We further conducted independent DEG analyses among the PFC, EC, and SFG regions to obtain reliable benchmark results. As shown in Figure 3C-F, the number of DEGs in multiple cell types is still higher than cell-type-specific DEGs when we dismissed the regulated directions. In contrast, our results revealed that both up- and down-regulated DEGs were highly cell-type specific (Additional file 2: Figure S3), consistent with a previous report [13].

**Figure 3.**
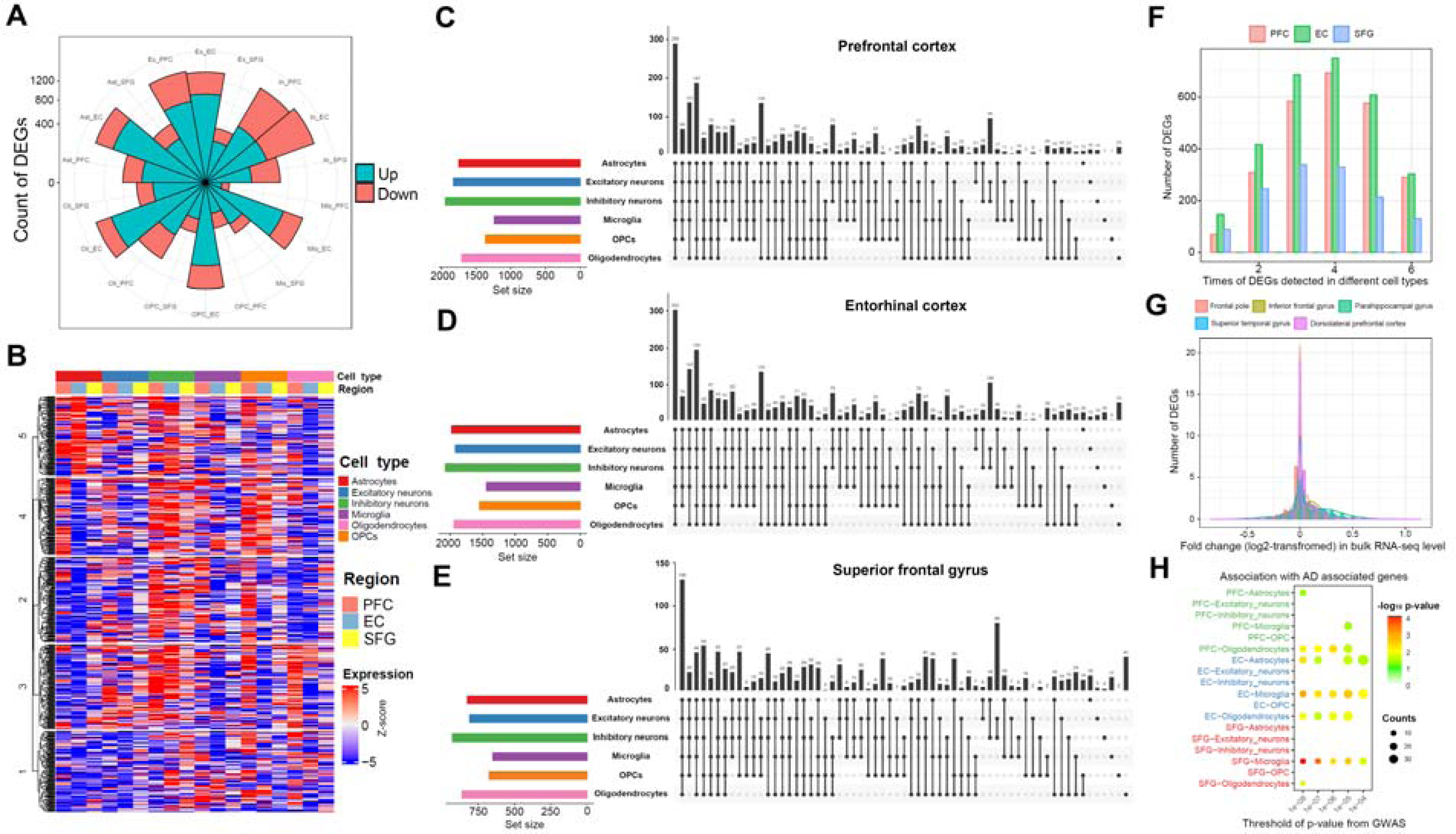
Comparison of differentially expressed genes among cell types and regions. **(A)** Summary of number of DEGs over various cell types and regions. **(B)** Heat map of 532 conserved DEGs (consistent regulatory direction in three brain regions in at least one cell type) over different cell types. **(C-E)** UpSetR plot shows the number of common or cell type-specific DEGs among **(C)** PFC, **(D)** EC, and **(E)** SFG, without taking regulation direction into consideration. **(F)** Summary of DEG frequency across six major cell types indicating the majority of them were differentially expressed in multiple cell types. **(G)** Comparison of DEGs from AD scRNA-seq/snRNA-seq and two bulk AD cohorts (MSBB and ROSMAP). **(H)** Association of DEGs (snRNA-seq derived) and AD-associated genes based on genome-wide association studies.

We also conducted DEG analysis using bulk RNA-seq data from MSBB-AD [16] and the ROSMAP [17, 18] cohorts across five brain regions. As expected, due to a large proportion of DEGs with opposite direction in different cell types, most DEGs detected at the single-cell level did not show significant differences at the bulk level (Figure 3G). For example, one previous study observed that *APOE* was up-regulated in microglia but downregulated in astrocytes [13]. To understand in which cell type DEGs were most vulnerable to AD dysregulation, we subsequently linked them with GWAS summary statistics [27] to evaluate the association between cell-type-specific DEGs and AD susceptible genes. As shown in Figure 3H, AD susceptible genes were mainly enriched in glial DEGs rather than in neuronal cells. We further conducted tissue-specific enrichment analysis and temporal-specific analysis using software *deTS* [31] and our previous imputed BrainSpan data [5]. As shown in Additional file 2: Figure S4, these DEGs were mainly enriched in adipose or immune-related organs, and they were significantly associated with late prenatal and early postnatal stages.

A common mechanism of gene regulation is the binding of TFs. Therefore, we used a deep learning-based model, DrivAER, to identify potential key TFs within different cell types of scRNA-seq. TF annotations were from MSigDB [23] and TRRUST [24]. Our analysis identified both common and cell type-specific TFs (Additional file 2: Figure S5). Recently, a network approach combined with scRNA-seq has been proposed as a valuable strategy to investigate molecular mechanisms in AD [10]. Therefore, we reconstructed the regulatory network by integrating the conserved DEGs. As shown in Figure 4A, 107 genes were embedded in the AD-associated regulatory network. We grouped them into six functional categories: transporter, cell differentiation, cell adhesion, energy metabolism, immune response, and neuron development (Additional file 1: Table S5). Those genes had substantial functional discrepancies among different cell types and directions. For example, the up-regulated genes of inhibitory neurons were mainly enriched in cell differentiation, cell adhesion, and energy metabolism. In contrast, down-regulated genes of glia cells were enriched in transporter and neuron development (Figure 4B).

**Figure 4.**
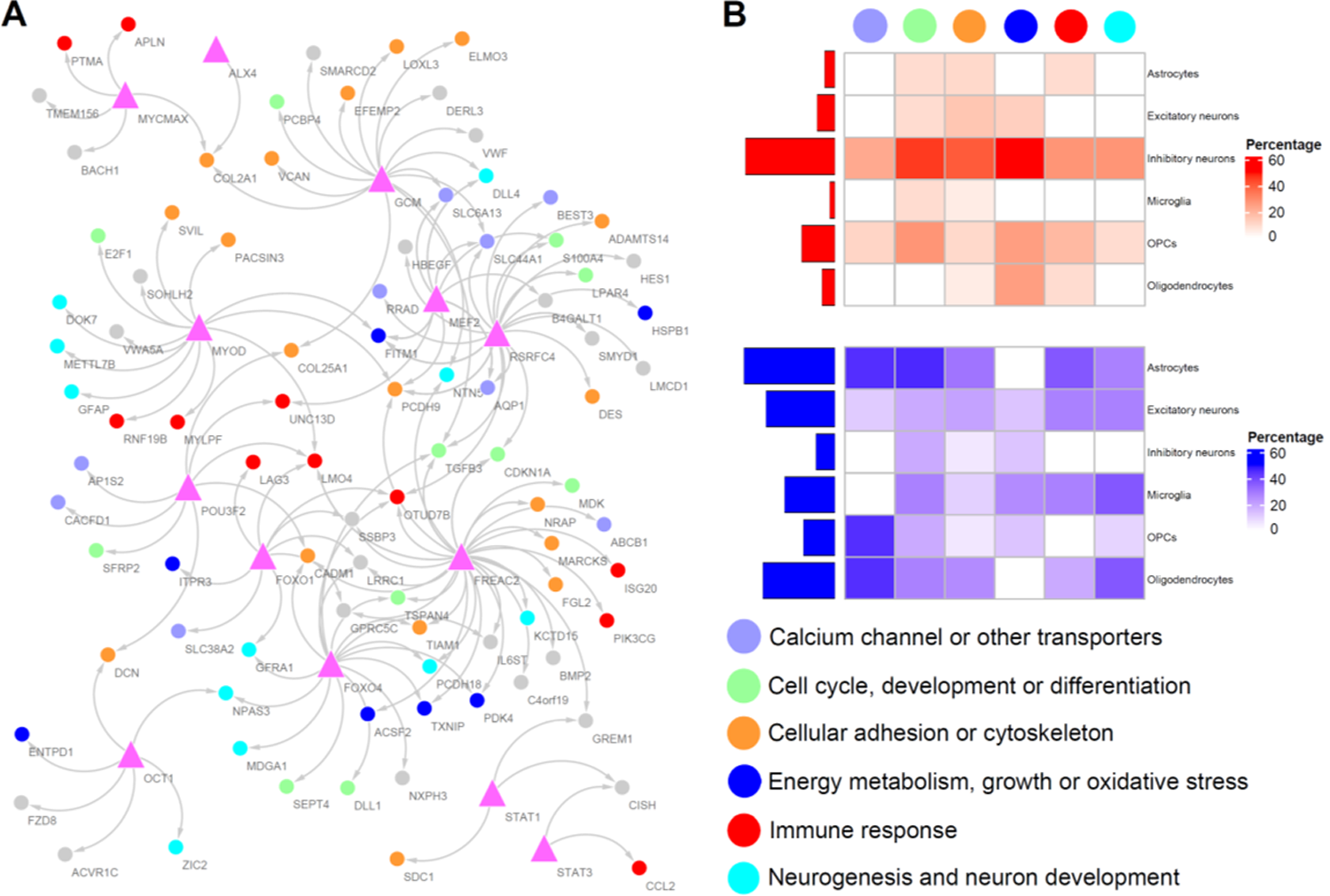
DrivAER unveiled key transcription factors and molecular networks among AD-associated cell types. **(A)** Visualization of interplays between TFs and DEGs from GSEA C3 collection: regulatory target gene sets and TRRUST databases. Triangle and circle represent transcription factor and DEG, respectively. **(B)** The distribution and overlapping percentage of up or down-regulated gene pairs among different functional categories.

### AD-associated cells were individual-specific: subpopulation analysis

To dissect cellular heterogeneity, we further performed unsupervised clustering for each of the major cell types. We identified a total of 54 cell subtypes with a median of 824 cells in each subtype (range between 196 to 31,649). Each subtype was composed of at least eight cell subtypes (median 33, Figure 5A). Unsurprisingly, the number of subtypes did not show any significant difference between AD case and healthy control among the three brain regions (*p*-value > 0.05, Wilcoxon-sum test). However, we observed a slight difference in cell type enrichment between the EC and SFG (Figure 5B). We further examined their relative composition to pathological features and brain regions (PFC, SFG, or EC). All cell subtypes were composed of at least three different samples (median 31, Figure 5C). In addition, we observed an enrichment of the cellular population in AD brain for most cell types. For example, the cellular subpopulations of Ast1, Ast2, Oli3, Oli4, Oli5 were enriched in the AD population. In contrast, Ast6, Ex3, Ex6, In1, In8, In9 were depleted in AD (FDR < 0.001, hypergeometric test). In addition, we also observed an overrepresentation of cellular subtypes in specific brain regions (Figure 5C). Most of these patterns were robust and did not change when other unsupervised clustering methods were applied (Additional file 2: Figure S6).

**Figure 5.**
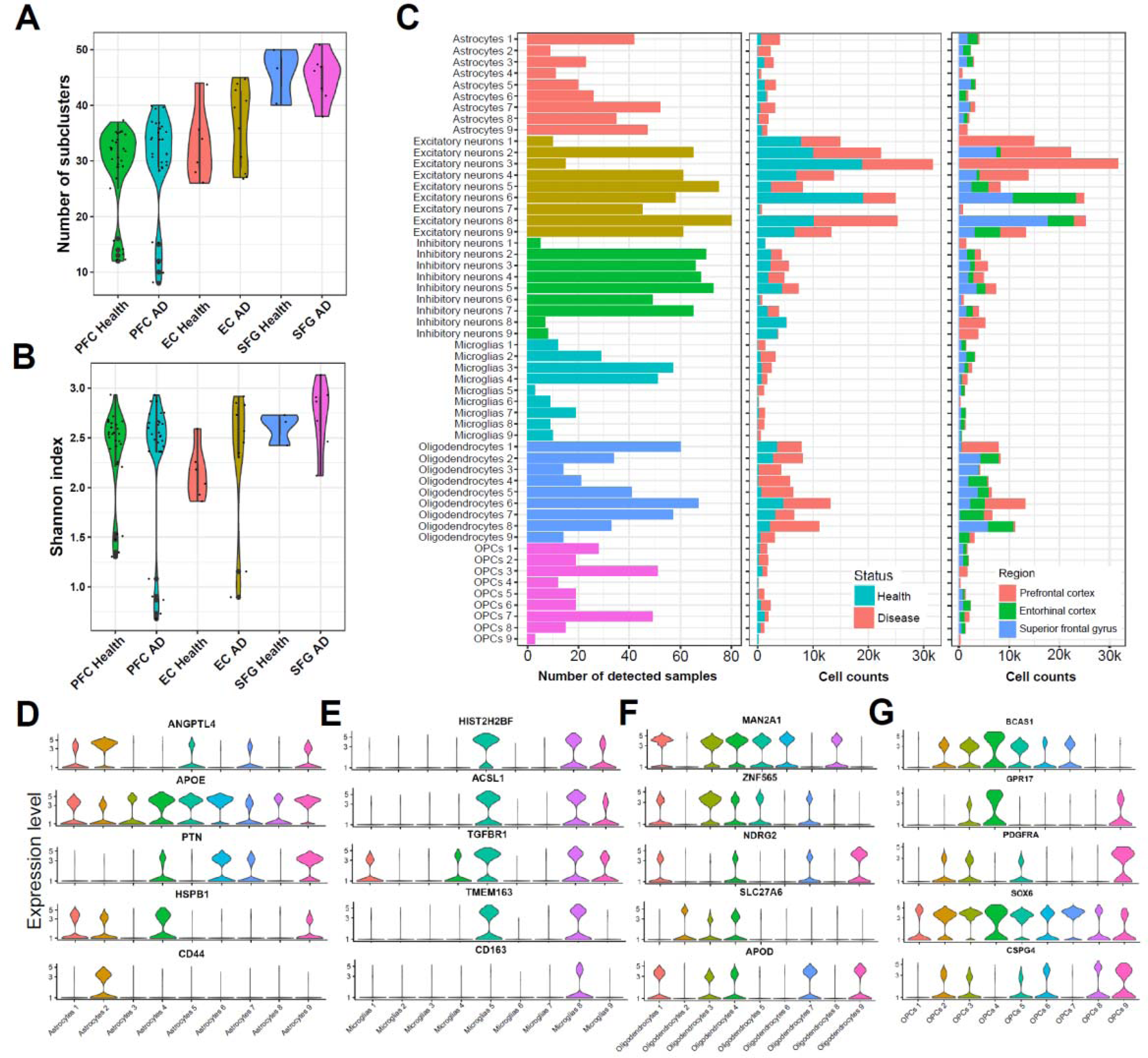
Diverse cell subpopulations in Alzheimer’s brain. **(A)** Richness and **(B)** α-diversity (Shannon index) comparison between control and disease groups at the gene level. Violin plots showed the richness without significant difference in all regions, while the α-diversity value s elevated in both the EC and the SFG. **(C)** Distribution analysis of different cell subpopulations among different individuals (lower frequency, higher specificity), status (AD enriched or depleted), and regions. **(D-G)** Violin plots of representative marker gene expression levels among different glia subpopulations, including **(D)** astrocytes, **(E)** microglia, **(F)** oligodendrocytes, and **(G)** OPCs.

Accumulating evidence has highlighted the importance of glia change in AD [10]. In astrocytes, we noticed that *ANGPTL4*, a gene encoding hypoxia-induced factor that is the target of peroxisome proliferator-activated receptors (PPARs) and regulates lipid metabolism, is significantly increased in the Ast2 cluster (Figure 5D). This result suggested vascular and lipid dysfunctions, which is in contrast with the finding from a previous study with participants with subjective memory complaints [39]. We also observed an increase of *CD44* gene expression in Ast2. Recent evidence shows that high expression of *CD44* is associated with long, unbranched processes and “fibrous”-like astrocytes, whereas the “protoplasmic” astrocytes, which display a “bushy” morphology, exhibit no *CD44* expression [40]. In contrast, *APOE* was downregulated in Ast2 [41, 42]. In microglia (Figure 5E), some subtypes exhibited higher levels of *TGFBR1*, which has been implicated in the regulation of functional aspects of several distinct immune cell populations in the central nervous system [43, 44], such as insulin regulation, with higher levels being associated with decreased glucose uptake [45]. In addition, we observed *TMEM163,* which modulates the purinergic *P2X* receptor activity, and thus purinergic signaling [46], by modulating *P2XR* channel properties [47], and that express glutamatergic and γ-aminobutyric acid receptors [48]. We also reported in a previous study that the macrophages marker *CD163* showed a strong expression level in the plaque-associated microglia subpopulation [49]. Furthermore, the strong AD-associated immunomodulatory gene *MAN2A1* [33] is increased in most AD-associated oligodendrocyte populations (Figure 5F). Recent evidence has demonstrated that increased *NDRG2* is colocalized with NFTs and senile plaques in sporadic AD patients [50]. Collectively, we noticed that most subpopulations of the neuronal or glial cell were detected in less than ten samples, suggesting that they contribute distinctively to different individuals with AD.

### More diverse cell-cell communication patterns in AD brain

As the most complex organ of the human body, brain development involves a finely-tuned cascade of interactive events [5, 51]. Although previous studies had investigated the DEGs and enriched pathways among different AD cohorts [12–15], their respective cellular interactions (L-R pattern) have not yet been rigorously characterized [52]. Given the discrepancy regulation patterns among different cell types (Figure 3B), we sought to conduct L-R communication analysis in a cell subpopulation manner to determine if there are any unexpected intercellular communication patterns in AD brains. Given our observation of slight batch effects after integration, we first conducted an ensemble analysis of the above 54 cell sub-populations. As shown in Additional file 2: Figure S7, we present the top L-R incoming and outgoing pathways estimated by CellChat [25]. Unsurprisingly, the snRNA-seq offers substantial advantages in detecting cellular and transcriptional diversity in multiple human tissues [53, 54]. We found some AD-specific or healthy brain-specific pathways, *e.g.*, secreted phosphoprotein 1 (*SPP1*) (*SPP1* - *CD44*), suggesting positive regulation of immune cell activation and infiltration [55], in astrocytes subpopulation 2 (Ast2) (Additional file 2: Figures S8A); *VISFATIN* (*NAMPT* - *INSR*), which is a pro-inflammatory adipokine up-regulated by inflammation [56] that can induce sickness response in the brain [57] and consequent anorexia and weight loss (Additional file 2: Figures S8B) in Ast2; Semaphorin 4 (*SEMA4*) (*SEMA4D* – *PLXNB1*), which is implicated in the control of cell migration and neuronal connectivity [58, 59], in Ast2_in_, Ast7_in_, and Ast8_in_.

We further investigated their cellular interactions in different brain regions to discern the conserved and region-specific dysregulated communication patterns in AD. The overall number and the strength of cellular interaction were increased in all three AD brain regions (PFC, SFG, and EC) compared to healthy control (Figure 6A), indicating a substantial response to stress and stimuli in AD. As shown in Figure 6B, we observed that OPC was the most active cell type, while microglia was the least active. This might be due to the low sensitivity of snRNA-seq to detect microglial activation genes in the human brain [60]. On the other hand, our cellular communication analysis showed that the most robust differential interactions were detected among inhibitory neurons, OPCs, and oligodendrocytes (Figure 6C). Next, we further examined the information flow for each signaling pathway, which led to a deeper understanding of the conservation and context-specific communicating patterns in the brain of individuals with AD. As shown in Figure 6D, most of the dysregulated signaling pathways involved in neurogenesis and cell-cell adhesion were significantly increased in all those three brain regions. For example, *NRXN* (neurexin) was involved in synaptic formation, elimination, plasticity, and maturation [61]; *NRG1* (neuregulin 1) influenced Aβ load, synaptic integrity, neuroprotection, and influence cognitive function in AD [62]; *PTPRM* was associated with axon guidance and cell proliferation, migration, and invasion [63]. We presented the communication probabilities and contribution of some representative L-R pairs in Additional file 2: Figures S8 and S9. Collectively, the overall communication probabilities changes between AD and control showed a higher overlap degree among different regions.

**Figure 6.**
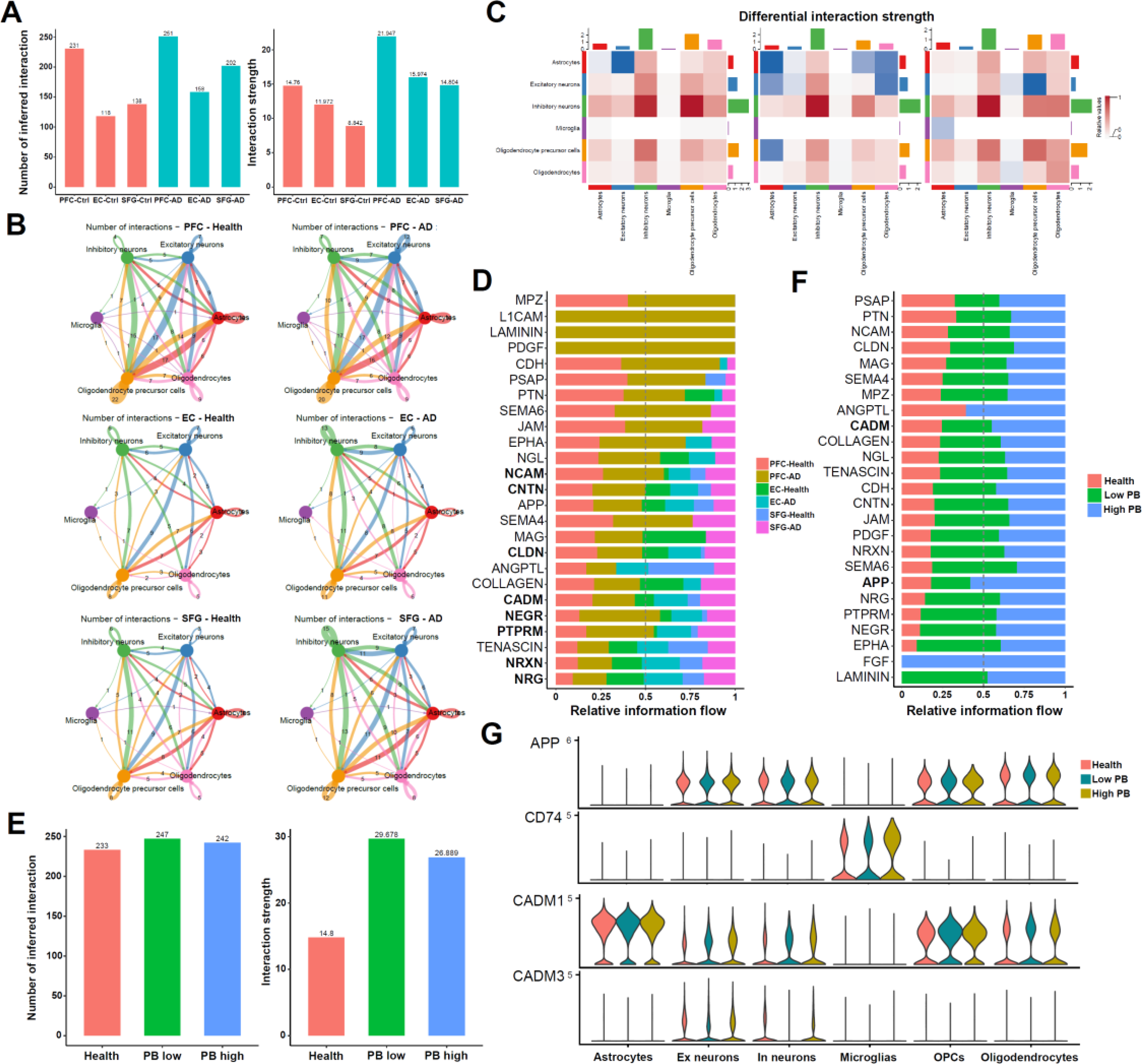
Cell-cell communications among six major brain cell types. **(A)** Bar plot showing the total number of interactions and interaction strength of the inferred cell-cell communication networks between AD brain and healthy control. **(B)** Circle plots summarizing the number of interactions among different cell types in each brain area. The thickness of the lines connecting cells indicates the interaction strength. **(C)** Heat map showing the overall information flow of AD differential interactions among individual cell types in each brain area. **(D)** Top representative differentially expressed L-R signal pathways between the brains of healthy controls and AD participants. **(E)** Bar plot showing the total number of interactions and interaction strength of the inferred cell-cell communication networks among healthy control, low and high pathology burden individuals. **(F)** Top representative differentially expressed L-R signal pathway among healthy control, low, and high PB groups. **(G)** Violin plots showing two representative L-R signal pathways (*APP* and *CADM*) in the high PB group. L-R: ligand-receptor. PB: pathology burden.

The neuropathological manifestations of AD start years before the development of any visibly apparent cognitive symptoms [4]. Therefore, whether the communication patterns in patients with more severe AD-associated dementia are divergent or conserved with early-stage AD remains to be verified. As shown in Figure 6E, the overall number and the strength of cellular interactions in the low PB group is even higher than the high PB group, which may imply divergence in the early stage of AD. In addition, we observed high degrees of overlap between high PB and low PB (vs. healthy control, as shown in Figure 6F). These findings may indicate that major transcriptional changes take place before the development of severe pathological alterations [13]. Of interest, we observed that the signaling pathway of APP-CD74 was enhanced in high PB but not in low PB individuals (Figure 6F). We provided their expression level in *CD74*, which encodes an antigen-presenting protein that can interact with mature *APP* to reduce Aβ production and thereby improve AD-associated memory deficits [64]. This finding is consistent with previously observed *CD74* expressed in NFTs AD brain [65]. Interestingly, another signaling pathway, cell adhesion molecule (*CADM1*), was up-regulated in neuronal cells of high PB patients; this cell adhesion molecule also has a pivotal role in the pathogenesis, severity and progression of AD from the perspective of neuro-inflammation, Aβ metabolism, cell plasticity and vascular changes [66].

### Prediction of AD brains on bulk RNA-seq data by using snRNA-seq markers

To illustrate the diagnostic value of AD-associated markers by scRNA-seq/snRNA-seq, we utilized the conserved DEGs and dysregulated L-R genes and constructed a series of random forest classifiers to detect the AD and MCI brains from bulk level (from MSBB-AD [16] and ROSMAP cohorts [17, 18]). First, we randomly selected 90% of the samples in the MSBB-AD dataset for training, leaving the 10% remaining for testing. Ten repeats of 10-fold cross-validation (total 100 tests) in the training set consisting of 118 health and 600 AD brains, or 126 MCI brains, led to the optimal selection of 15 and 40 markers that performed well on the training and test sets, respectively. We used the AUROC to evaluate the performance of the classifiers. By this measurement, the control-AD classifier achieved an AUROC value of 76.9% and 80.43% (Figure 7A), while the control-MCI classifier achieved an AUROC value of 79.8% and 85.4% on training and test (Figure 7B), respectively. However, when we did not include dysregulated L-R features in those models, the AUROCs were 2%-4% lower (Additional file 1: Table S6). Therefore, a model which could discover the cell-cell internal interaction would be more appropriate to obtain a complete picture of cellular response in AD brains under bulk RNA-seq level.

**Figure 7.**
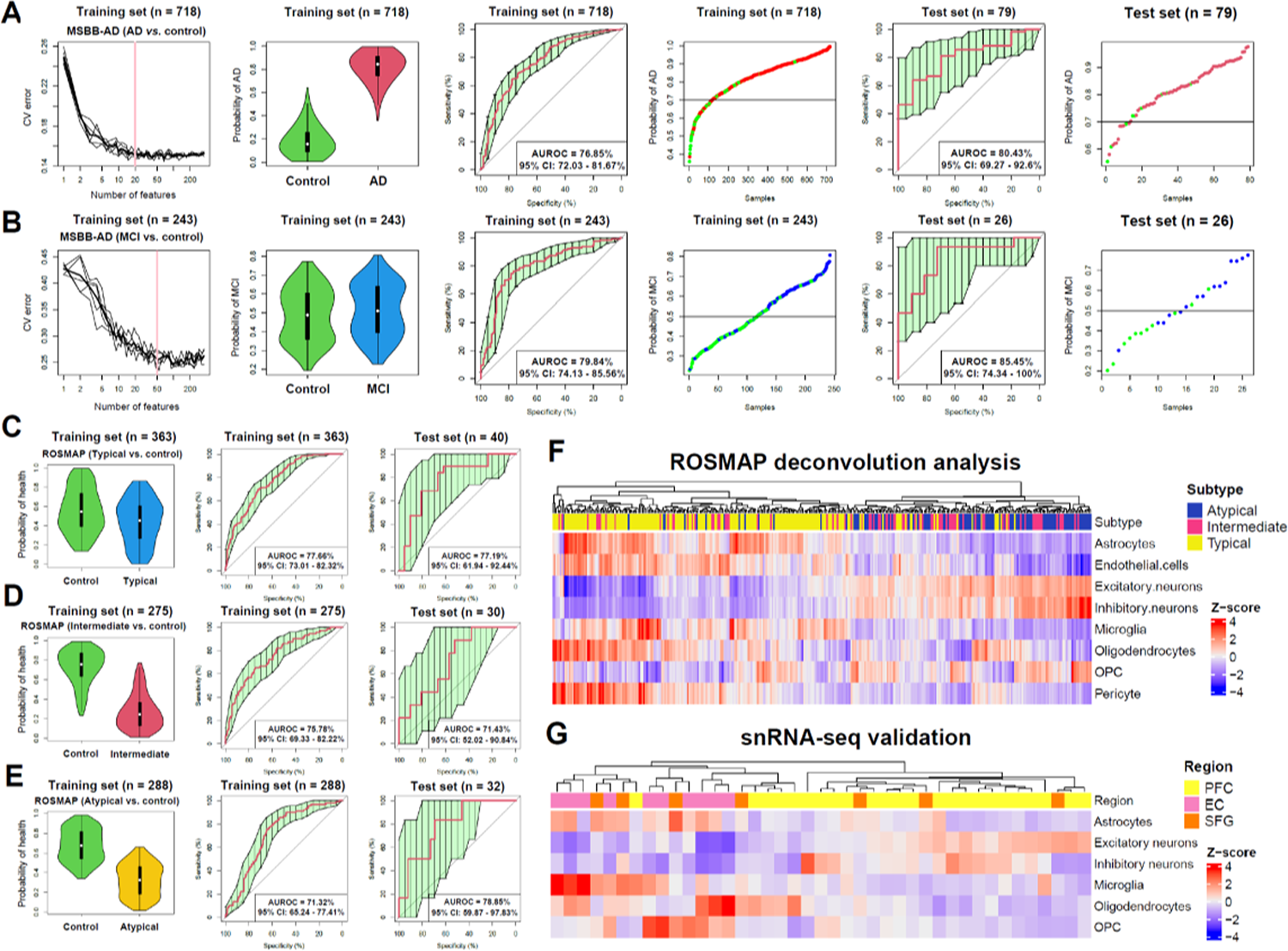
Identification of MCI and AD brains from healthy control in the MSBB and ROSMAP cohorts at bulk RNA-seq level. **(A)** The models were trained (using 572 conserved DEGs) between the healthy controls and AD samples (n = 118 and 600), or **(B)** healthy controls and MCI samples (n = 118 and 126) on the MSBB cohort. From left to right, they represent (1) the distribution of ten trials of 10-fold cross-validation error in random forest classification models. The bold black curve indicates the average of the ten trials. The pink line marks the number of gene features in the optimal set. (2) Violin plot for the probability of AD or MCL in the cross-validation training set according to the model in (1). (3) The AUROC for the training data sets, along with a 95% confidence interval. (4) Classification of the train set consisted of healthy control, AD, or MCI samples. (5) AUROC for the test sets, along with a 95% confidence interval. (6) Classification of the test set. **(C-E)** The models were trained between three AD groups: **(C)** typical, **(D)** intermediate, and **(E)** atypical subtype of AD and the healthy controls the ROSMAP cohort. From left to right, they represent (1) violin plot for the probability of AD in the cross-validation training set, (2) AUROC for the training, and (3) test data sets. **(F)** Hierarchical clustering of cell type deconvolution results of ROSMAP bulk cell RNA sequencing data. Because the imputed cell-type relative abundance differed in several scale orders, we took z-score as the normalized cell type relative abundance to elucidate the correlation between AD subtype and cell type relative abundance. **(G)** Hierarchical clustering of cell type composition (z-scaled) of AD brains detected by snRNA-seq. The AD subtype label at the top was derived from previous bulk data in the ROSMAP cohort.

Subsequently, both top-performing random forest models were selected for further feature characterization. Collectively, the correlation of feature importance (mean decrease accuracy) in the control–AD model was significantly correlated with the control–MCI model (PCC = 0.26, *p*-value = 4.82 × 10^−7^). GO and KEGG enrichment analysis demonstrated top genes enriched in MCH protein complex, ECM-receptor interaction, focal adhesion, cell adhesion molecules, and extracellular matrix structural constituent (Additional file 2: Figure S10). Among these features, *GFAP* and *CPVL* showed high importance scores in both AD and MCI classifiers. Some other features, including *JUNB*, *BYSL*, *SDS,* and *GNG11,* showed a strong importance score in the MCI classifier but not in the AD classifier. Nevertheless, given the heterogeneity and complexity of the AD phenotype, a recent study classified AD into three subtypes, i.e., an Aβ-predominant typical class, a tau-predominant intermediate class, and an atypical class [4]. Here, we investigated classifiers and possible drugs for repurposing using these previously defined in the literature subtypes.

### Prediction of AD brains across different subtypes

Based on the AD subtype information from a recent study (Additional file 1: Table S7) [4], we constructed three independent random forest models to investigate their unique molecular features from the ROSMAP dataset. As shown in Figure 7C, one model reached an AUROC of 73.01%-82.32% (95% CI) for the typical AD subtype, suggesting that the expression pattern in the typical subtype is more similar to snRNA-seq patterns. However, other two models only achieved an AUROC of 69.33% to 82.22% and 65.24% to 77.41% (95% confidence interval, CI) for the intermediate and the atypical AD subtypes (Figure 7D and 7E). Next, we investigated a comprehensive feature importance comparison among three independent models. The results revealed no significant correlation (*p*-value > 0.05) among the features of typical, intermediate and atypical subtypes (Additional file 2: Figure S10), suggesting distinct intrinsic molecular mechanisms [4].

By integrating AD subtype information with cell-type deconvolution results (Figure 7F), we uncovered that the cell type composition of neurons and OPCs decreased while astrocytes, endothelial cells, and microglia increased in the typical class of AD. On the other hand, the cell type composition of neurons and OPCs strongly increased, along with a decrease in different cell types in the atypical class [4]. To test the above hypothesis, we further investigated the relative cellular abundance of AD participants’ snRNA-seq data from the same ROSMAP cohort (Additional file 1: Table S8). As shown in Figure 7G, we observed a consistent downward trend of neuronal cell patterns in the typical type. In contrast, we observed that the relative composition of excitatory or inhibitory neurons in each half of the participants was strongly increased in the atypical type. Consistent with previous deconvolution results [4], we noticed these subtypes were neither determined by a high level of Aβ and tau accumulation, nor by differences in APOE risk allele genotype [41, 42]. Moreover, the end effects of neurodegeneration and inflammation from “typical” subtypes might show “opposite” molecular gene regulation in the “atypical” or “intermediate” types.

### Drug response analysis discovers repurposable and personalized drugs

Given the complexity and heterogeneity of AD, it can be challenging to identify drugs that benefit AD individuals universally. Therefore, we extended our previous method, called CeDR Atlas [35], to fine map drug response at cellular resolution and shed light on the design of both repurposable and personalized drugs at cellular level (Additional file 1: Table S9). To minimize the impact from batch effects, we focused only on the PFC region of 24 AD participants [13]. Next, we presented a patient–drug associated network to highlight significant associations (predicted by at least two among top 50 candidates). As shown in Figure 8A, we observed several potentially repurposable drugs, such as trichostatin, which demonstrated its broad applicability in 24/24 patients, acting especially in microglia, OPCs and inhibitory neurons. Recent studies report that trichostatin, an antifungal antibiotic that act as a histone deacetylase inhibitor, can effectively decrease neuroinflammation [67] in AD, and might help clear Aβ from the brain [68]. Histone deacetylases catalyze the removal of acetyl groups from histones and non-histone proteins and are epigenetic regulators; thus, inhibitors of histone deacetylases can increase acetylation and, consequently, gene transcription [69], preventing neurodegeneration. Histone deacetylases inhibitors have also demonstrated neuroprotective effects [70]. As such, trichostatin can ameliorate short-term recognition memory and long-term spatial memory in mice [71].

**Figure 8.**
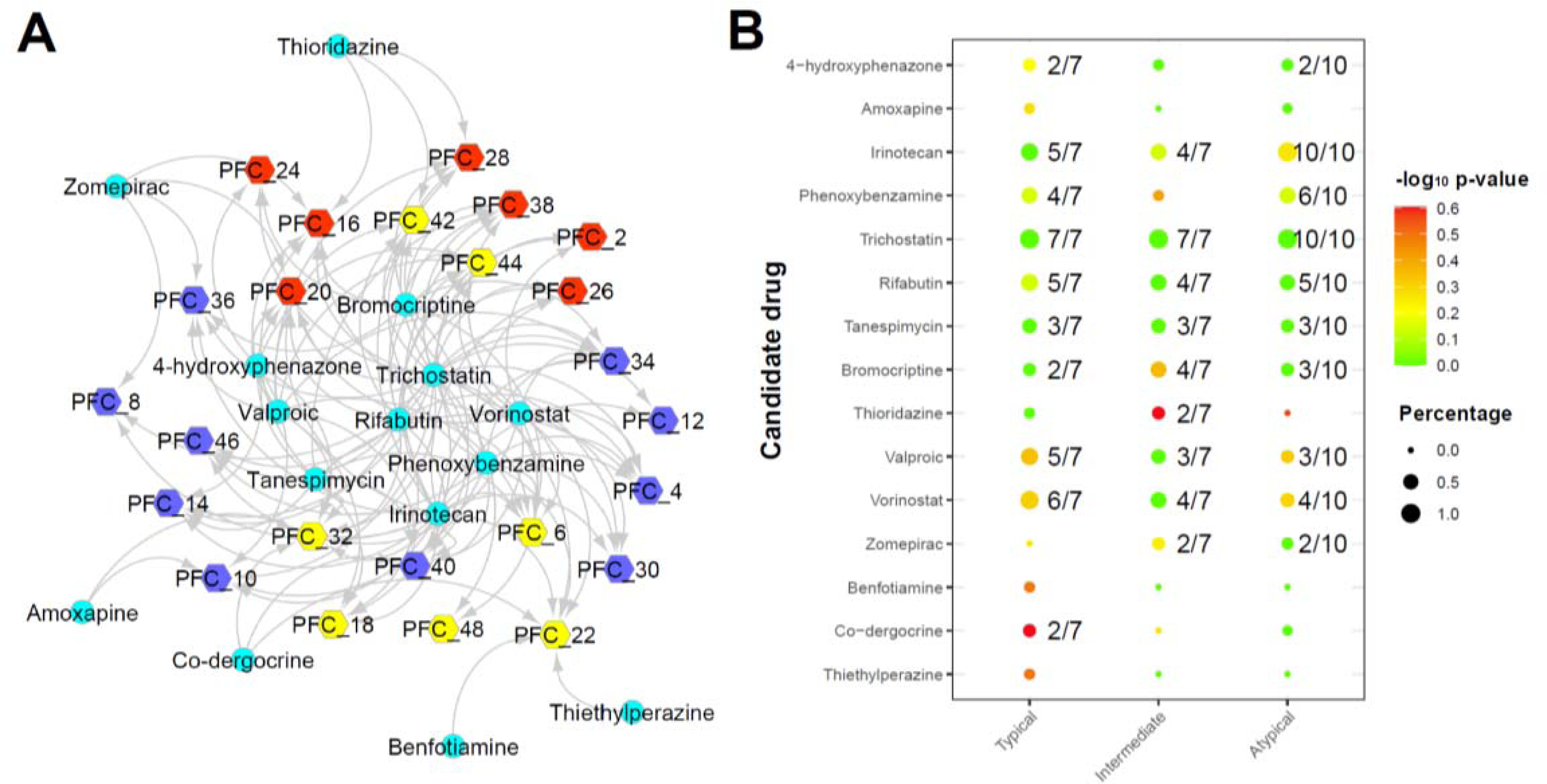
Drug repurposing analysis based on CeDR Atlas. **(A)** Network visualization of different AD individuals and top candidate drugs. Hexagon and circle represent AD individual and drug, respectively. Hexagons in yellow, red and blue color indicate typical, intermediate and atypical subtypes of AD, respectively. The network only shows the high confidence relationship which is predicted by at least two among top 50 drug candidates for each patient. **(B)** Association between different drug candidates versus three different AD subtypes. The size and color scheme reflect the percentage and −log_10_ (transformed *p*-value), respectively. Non-significant associations (less than 2 patients) were replaced by blank color.

Drug candidates [72] demonstrated distinct patterns across the three subtypes of AD. Vorinostat (14/24), an antineoplastic drug and also an histone deacetylase inhibitor [73] that induces hyperacetylation of α-tubulin and histones H3 and H4 [74], was effective in most of the typical subtype individuals (6/7, Figure 8B). The typical subtype is characterized by a decrease in neuronal cell populations. Vorinostat is capable of independently induce neuritogenesis, the formation of neurites by neuronal cells, a process essential to neuronal development and formation of synapses, through activation of the MAPK-ERK pathway [74]. Accordingly, one study showed that vorinostat might be a useful therapeutic to inhibit the negative regulatory effects on synaptic plasticity [75]. Another study in an Alzheimer’s mouse model reported a decrease in the expression of genes related to synaptic plasticity, which was partially restored after treatment with vorinostat [76]. On the other hand, thioridazine, a phenothiazine antipsychotic, was superior in two of seven intermediate subtype AD individuals. It is well known that thioridazine works to treat psychosis by blocking dopamine receptors. A recent study reported that thioridazine could induce autophagy and suppress glioblastoma tumorigenesis [77]. Similarly, bromocriptine, a dopamine agonist compound that has been reported to show *in vivo* anti-Aβ effects [78], demonstrated its better applicability in the intermediate subtype (4/7). Although recent studies have reported no significant correlation between AD subtype and amyloid content [17, 18], we observed the amyloid content of individuals with AD matched to bromocriptine has a slightly higher amyloid level than the average level. Lastly, another potential drug, tanespimycin, a Hsp90 inhibitor, was predicted to be beneficial in 9 of 24 individuals with AD. However, tanespimycin possesses poor blood-brain barrier penetrance, thus other Hsp90 inhibitors might be more appropriate [79]. Phenoxybenzamine was beneficial in 11 of 24 patients. This drug is a non-selective irreversible alpha-adrenoceptor antagonist and has been reported to ameliorate neuroinflammation and pathology in 5xFAD mice [80], but it did not show specific association with any three AD subtypes. Although most associations between AD subtypes were not significant by Fisher’s exact test, we believe a more precise AD stratification step would help future therapeutic exploration.

## Discussion

Single-cell RNA sequencing enables us to elucidate the cell type heterogeneity of the human brain at the single-cell level resolution [81, 82]. Although powerful, most single-cell studies of the AD brain only focus on a limited number of samples and very specific brain regions due to its high cost and moderate throughput [12–15]. Participants typically consist of a heterogeneous group of participants across many AD subtypes, and each subtype possesses distinct dysregulated pathways or a different combination of multiple dysregulated pathways [4]. Moreover, as pathological manifestations of AD start long before apparent cognitive symptoms, the AD dysregulated pathways do not affect all brain regions uniformly [6]. For instance, the hippocampal area demonstrates a greater subtyping signal than cortex regions [4]. In this study, to better understand the cellular, pathogenic, and regional heterogeneity in AD, we developed an approach to decipher the cell-specific mechanisms and to create an atlas of AD. We present an overview of our analytical approach in Figure 1. Firstly, we performed a comprehensive analysis of scRNA-seq/snRNA-seq datasets across three brain regions. Importantly, by doing this, we were able to gain several novel insights into cell subtypes, regarding both enrichment and depletion, with AD severity and brain region. By aggregating all of the scRNA-seq/snRNA-seq data and performing DEG analysis, we detected a number of DEGs across different brain regions for each cell type. Our results revealed that both up- and down-regulated DEGs were highly cell-type specific. Most conserved DEGs were regulated in opposite directions among different cell types. Secondly, we sought to determine if there were any unexpected conserved intercellular communication patterns in AD. After systematic identification of L-R interactions connecting cell surface proteins, we recognized novel mechanisms regarding intercellular communication in AD. We observed a list of conserved dysregulated communication patterns between AD and healthy control involved in neurogenesis and cell-cell adhesion in all three brain regions (Figure 6D). Moreover, we observed that most dysregulated patterns in the low PB group overlap with the high PB group, and significant increase of *APP* and *CADM* signal pathways from low to high PB (Figure 6F). These results implied analogous dysregulated communication patterns between MCI and AD. Comparable to the previous bulk sequencing studies [4], we developed a high-performance classifier to distinguish the MCI or AD samples from healthy control for each AD subtype. We noticed that models integrated with cell-cell communication features had better performance, implying the importance of intercellular communication involved in AD pathology. By applying CeDR for drug repurposing and subtyping analysis of AD, we detected not only that a histone deacetylase inhibitor, trichostatin, demonstrated broad applicability for all three subtypes of AD, but also some candidates, *e.g.*, vorinostat, another histone deacetylase inhibitor, would benefit more the specific AD subtype, and thioridazine and bromocriptine would be more useful to the intermediate subtype. To the best of our knowledge, personalized repurposing drug analyses on specific AD subtypes based on the combination of scRNA-seq/snRNA-seq has not yet been performed.

Our work has limitations. First, our study did not include a cohort of collected cells from different brain regions from the same participants. Compared to bulk cohorts (BrainSpan Atlas of the Developing Human Brain [83] and MSBB-AD [16]), the sample size and the collected regions remain relatively small (the hippocampal area is missing), which limited our ability to make entirely orthogonal comparisons. Second, the quality of the input data should be further improved in the future. For instance, one recent study demonstrated that snRNA-seq was insufficient to detect microglia activation genes in the human brain [60]. In addition, as transcript and protein abundance could be uncoupled by post-transcriptional processes, scRNA-seq/snRNA-seq data at the transcriptome level might not represent a fully accurate view of intercellular communication [25]. Therefore, emerging technologies such as single cell proteomics [84], cellular indexing of transcriptomes and epitopes by sequencing [85], nativeomics [86] and INs-seq [87] can further improve confidence [86]. Our findings from putative L-R networks were based on association, which warrants experimental validation in the future. Third, regarding target-based drug discovery, many efforts have been made on transcription-based drug discovery and implemented the concept of ‘anti-correlation between drug and disease signatures’ [88]. However, most drug profiles are available only on a restricted set of immortalized human cell lines [89]. To develop personalized treatments for each person with AD, a more comprehensive drug response profile should be collected or imputed [89]. We found trichostatin, a histone deacetylase inhibitor, to be effective across the three AD subtypes. However, vorinostat, another histone deacetylase inhibitor, appeared particularly beneficial in the typical AD subtype. The finding that these two different drugs with the same proposed mechanism of action appear to benefit different subtypes profiles might be a limitation of current drug repurposing analyses techniques. On the other hand, it may suggest the existence of a still unknown secondary mechanism of action of those drugs. Lastly, since the drug repurposing prediction is based on transcriptomic data of postmortem brain tissue, potential biomarkers for the specific AD subtype diagnosis that can be easily accessed, such as cerebrospinal fluid, blood [90], or non-invasive brain imaging, are highly desirable [4, 91].

In summary, we aggregated scRNA-seq/snRNA-seq data across the spectrum of AD brains for a comprehensive investigation. We observed strong cellular heterogeneity and transcriptional changes across different cell types in AD brains. Cell-cell communication analysis helped us to identify a core set of dysregulated cellular interactive events among the distinct AD subtypes, which can be utilized as important features to explain differences of prognosis and treatment response in AD. We characterize at least three different AD subtypes with unique cell-specific features. These results provide an avenue for AD stratification for future therapeutic exploration at the individual and cell type levels.

## Acknowledgements

GP and ZZ conceived the study. GP performed scRNA/snRNA-seq and bulk RNA-seq bioinformatics analysis and random forest model construction. YW, BSF, PJ, and AMM performed drug response analysis. BSF, AMM, and ZZ participated in results interpretation. GP, BSF, and ZZ wrote the manuscript. All authors read and approved the final manuscript. The authors thank the members of Bioinformatics and Systems Medicine Laboratory for valuable discussion, especially Drs. Hyun-Hwan Jeong and Yulin Dai and Mr. Andi Liu. We also thank Dr. Lukas Simon at Baylor College of Medicine for technical help. We acknowledge Dr. Bin Zhang at Icahn School of Medicine at Mount Sinai for insightful discussion on AD subtyping analysis.

## Code availability

The major steps for processing the scRNA-seq and bulk RNA-seq have been uploaded to <u>GitHub:</u> https://github.com/GuangshengPei/AD_scAtlas.
https://github.com/GuangshengPei/AD_scAtlas/blob/add-license-1/LICENSE.md

## Supplementary materials

### Additional file 1

Table S1. Summary of AD-related single-cell datasets used in this study.

Table S2. Data overview of scRNA-seq and snRNA-seq from scREAD database used in this study.

Table S3. MAGMA results of gene-based p-values based on recent meta-analysis of AD GWAS data.

Table S4. Conserved differentially expressed genes among six different cell types over three different brain regions.

Table S5. AD-associated regulatory network of key transcription factors and differentially expressed genes identified by DrivAER.

Table S6. Differential expressed ligand-receptor pairs between AD brains and healthy control.

Table S7. Clinical and AD subtyping information from ROSMAP cohort.

Table S8. Clinical and predicted AD subtyping information of 24 snRNA-seq datasets from prefrontal cortex region.

Table S9. Drug repurposing analysis results of 24 prefrontal cortex specimens based on CeDR Atlas algorithm.

### Additional file 2

**Figure S1.**
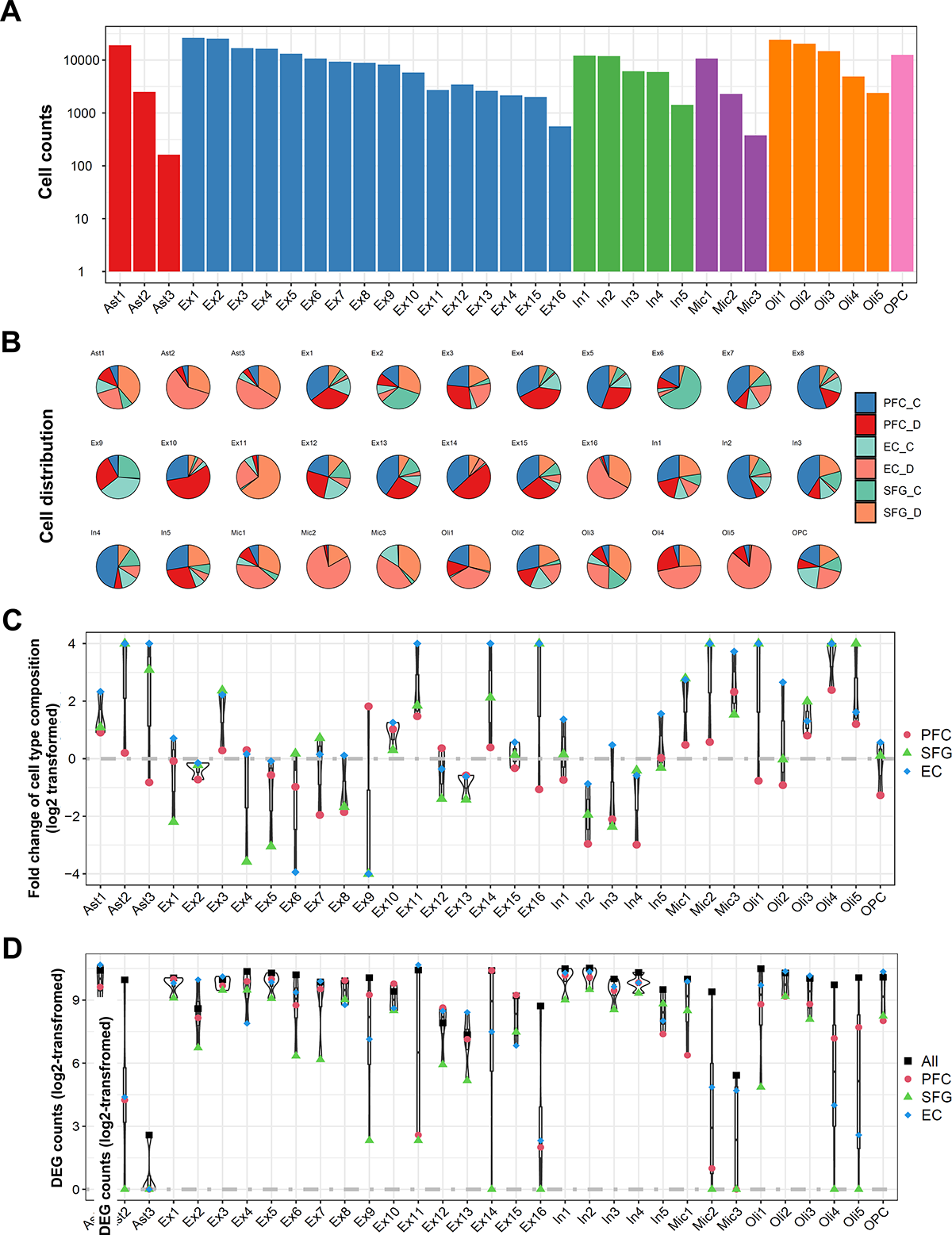
Overview of 33 distinct cell clusters in AD brains. **(A)** Cell counts. **(B)** Cell distribution. **(C)** Fold change of cell type composition. **(D)** DEG counts between AD brain and healthy control.

**Figure S2.**
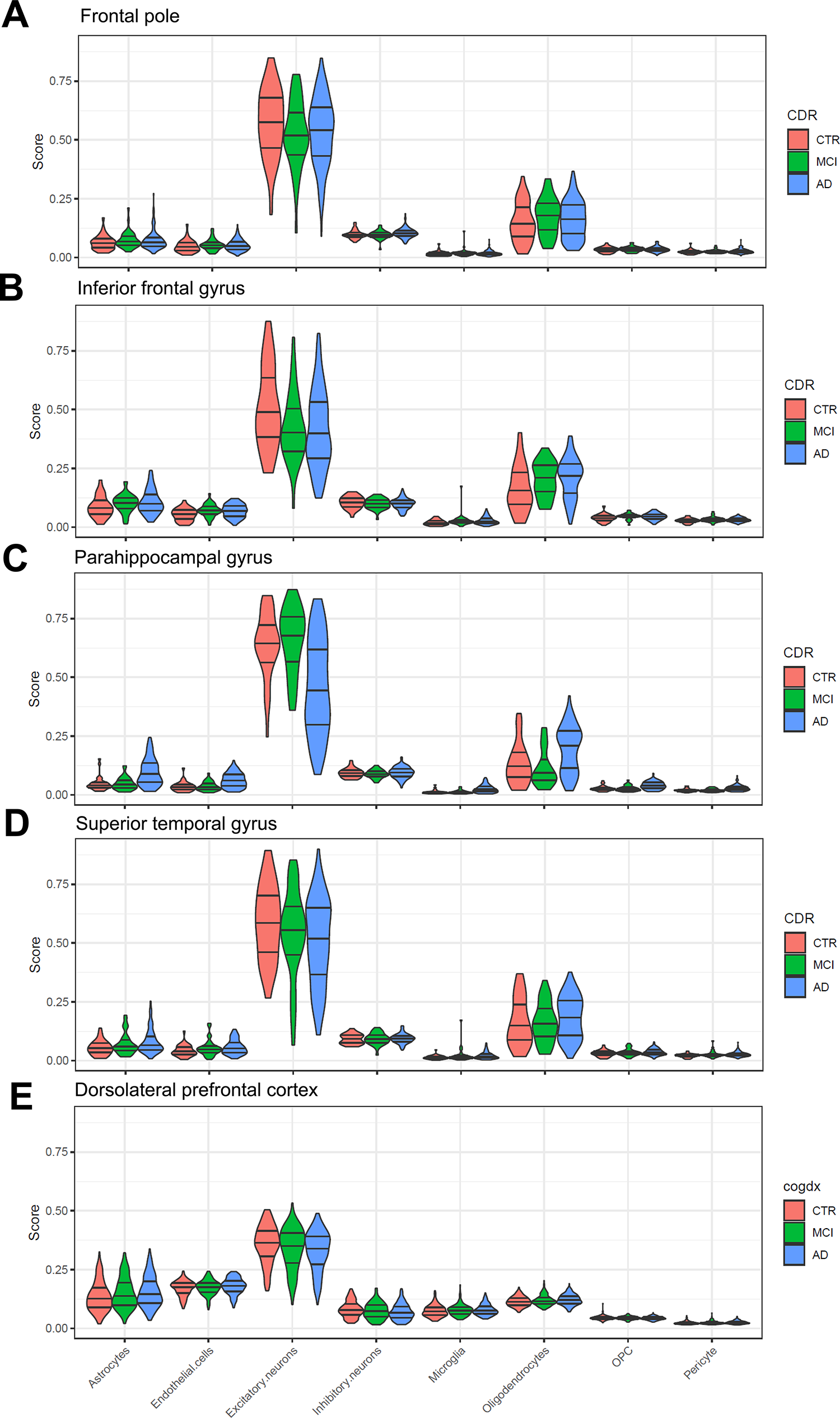
Change of different cell type compositions based on cell type deconvolution analysis on MSBB and ROSMAP cohorts. **(A)** Frontal pole. **(B)** Inferior frontal gyrus. **(C)** Parahippocampal gyrus. **(D)** Superior temporal gyrus. **(E)** Dorsolateral prefrontal cortex.

**Figure S3.**
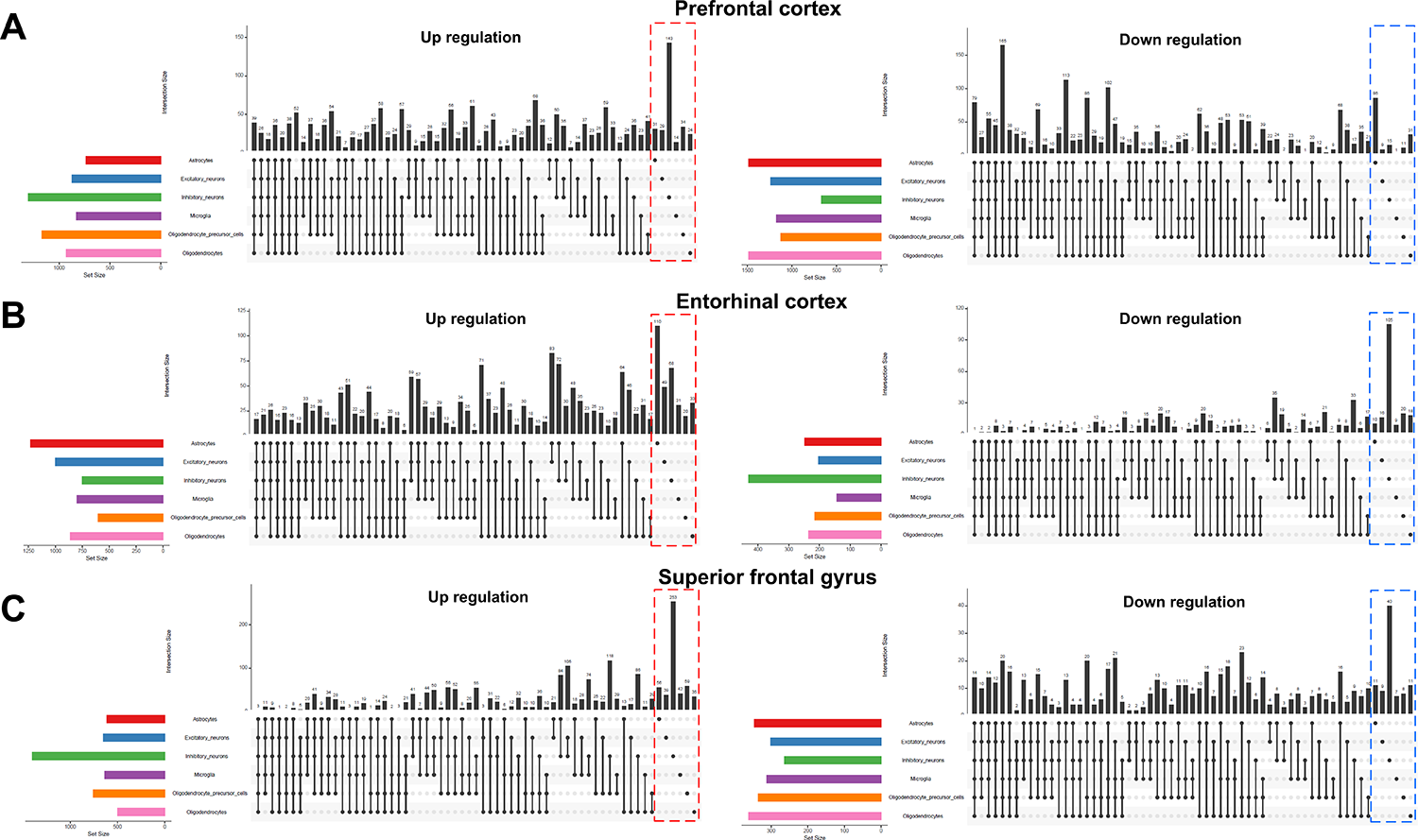
Differentially expressed genes comparison analysis among different cell types and regions. The number of common or cell type-specific up-regulated or down-regulated genes among **(A)** PFC, **(B)** EC, and **(C)** SFG were presented separately.

**Figure S4.**
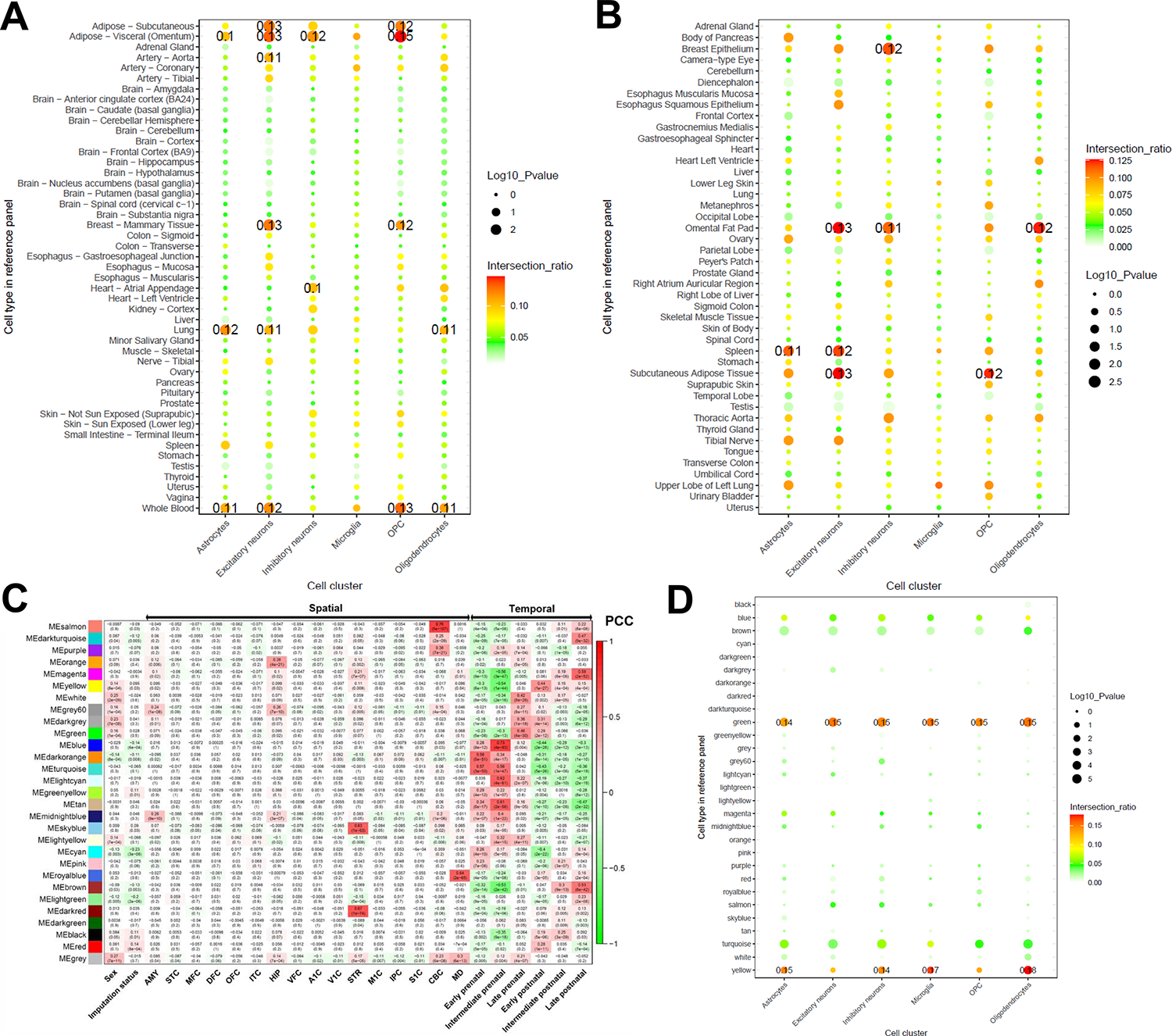
Tissue-specific enrichment analysis and temporal/spatial enrichment analysis of DEGs among different cell types. **(A)** GTEx and **(B)** ENCODE panels by *deTS* software. **(C)** WGCNA analysis results from BrainSpan and **(D)** module enrichment analysis among 29 WGCNA modules. The color and size reflect the shared gene counts and −log_10_ (transformed *p*-value).

**Figure S5.**
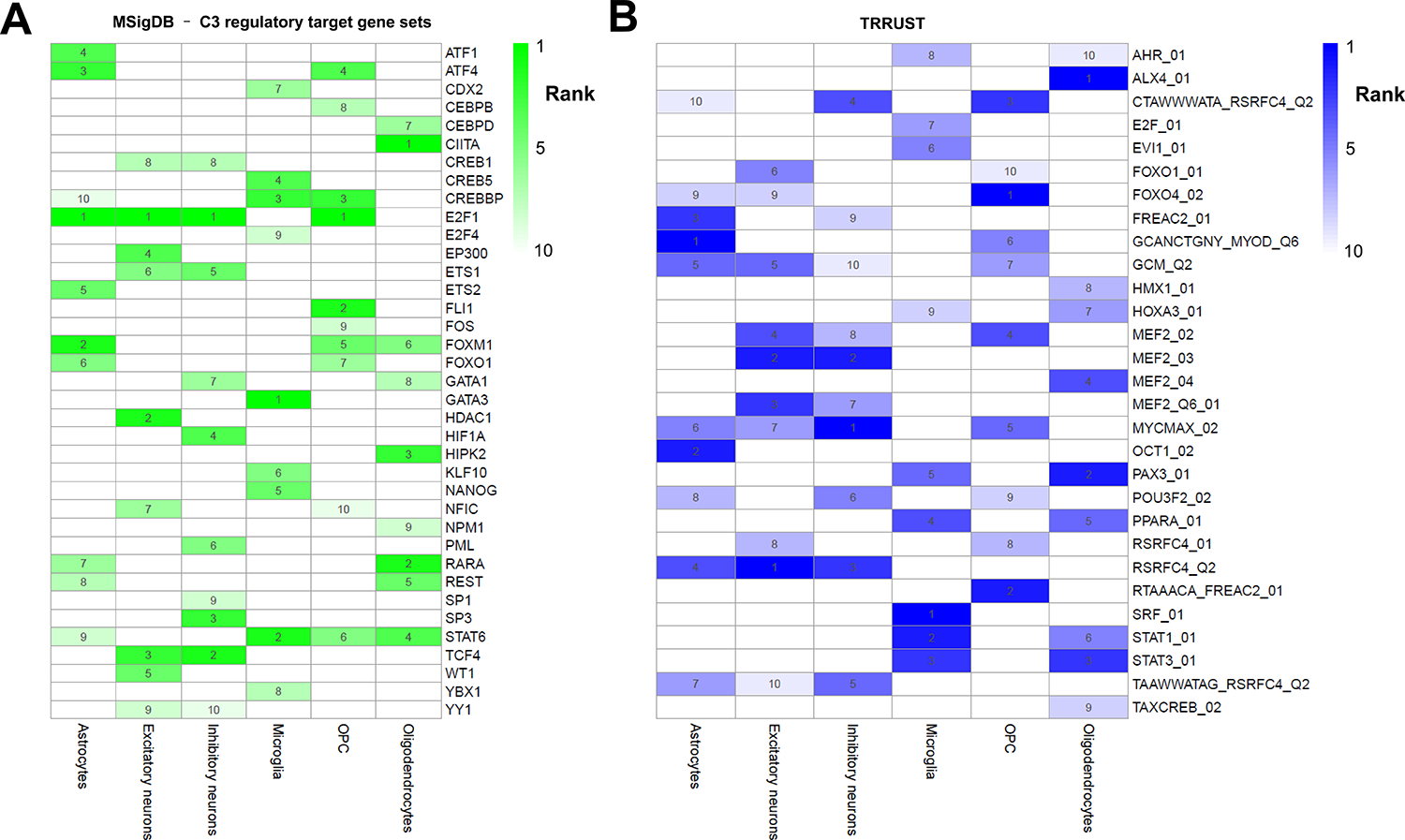
DrivAER analysis identified the top 10 most associated transcription factors differentially activated among different cell types. Transcription factor – target genes based on **(A)** GSEA C3 collection, or **(B)** TRRUST database.

**Figure S6.**
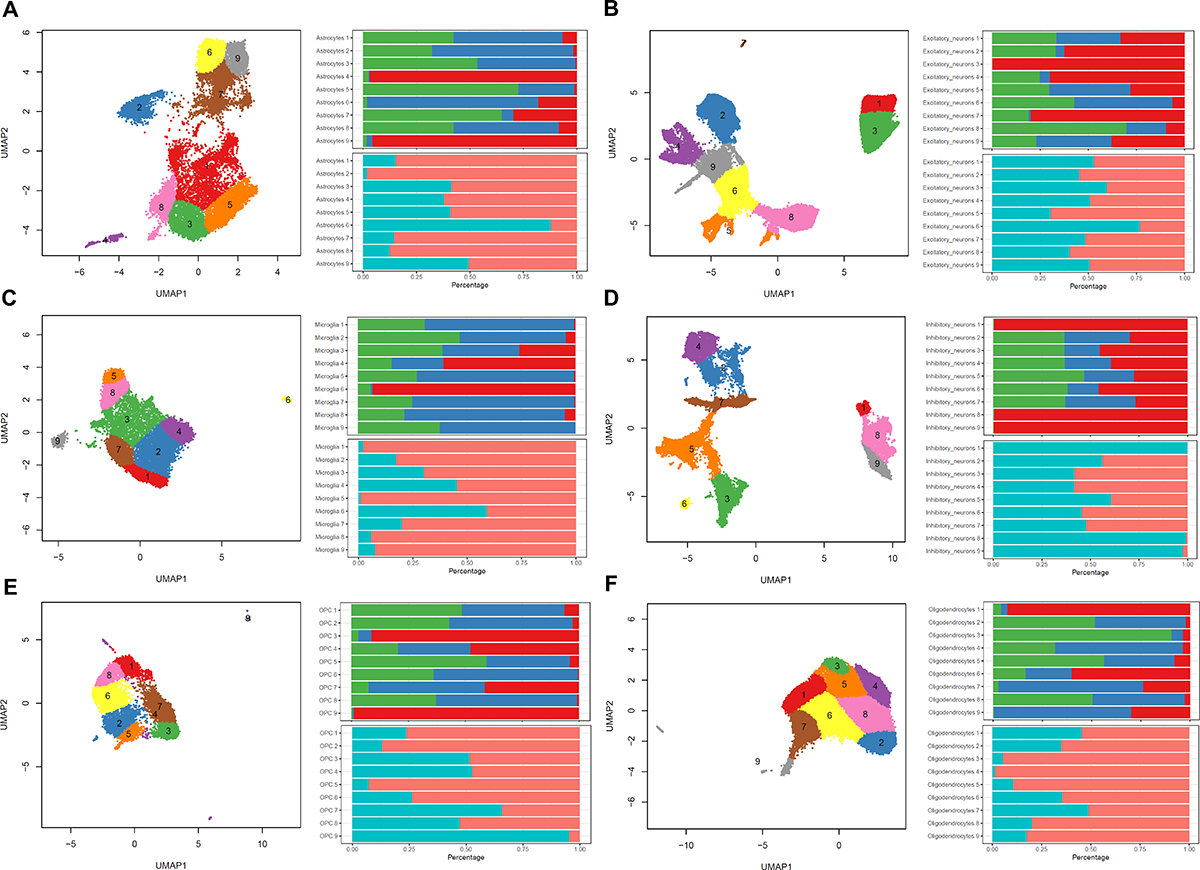
UMAP analysis and cell subpopulations analyses for different cell types by hierarchical clustering. **(A)** Astrocytes. **(B)** Excitatory neurons. **(C)** Microglia. **(D)** Inhibitory neurons. **(E)** OPCs. **(F)** Oligodendrocytes.

**Figure S7.**
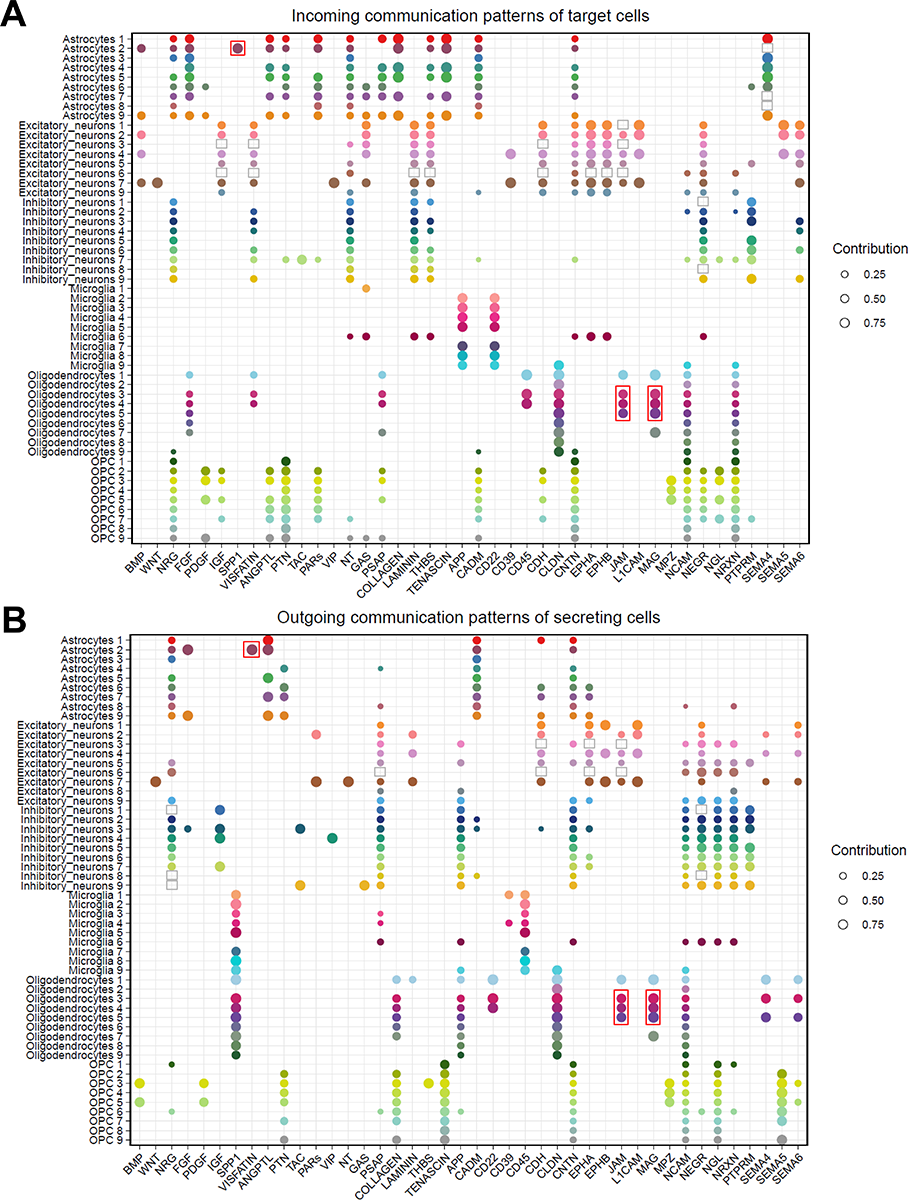
Overview of top ligand-receptor incoming and outgoing pathways among 54 cells subpopulations estimated by the CellChat software. **(A)** Incoming communication patterns. **(B)** Outgoing communication patterns.

**Figure S8.**
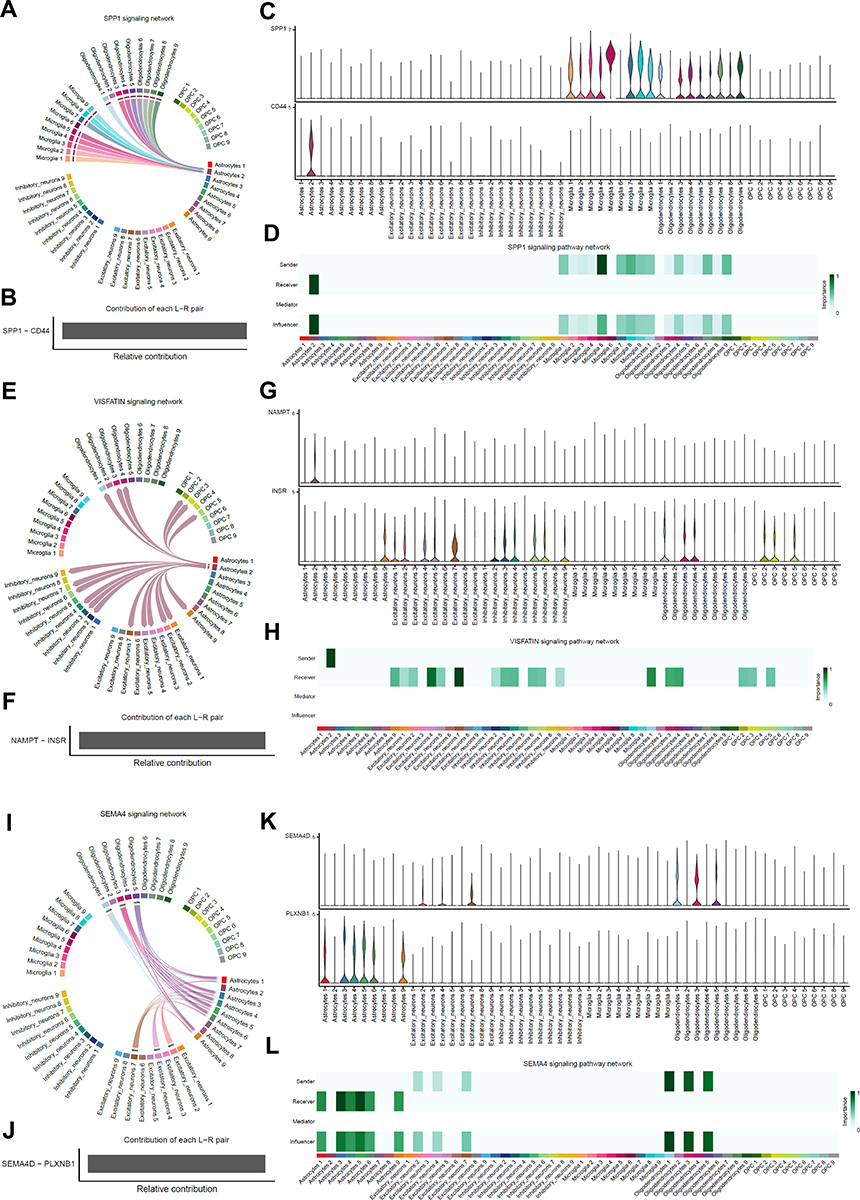
Three representative ligand-receptor signal pathways among 54 cell subpopulations. **(A)** Circle plot summarizing the interactions strength of the *SPP1* pathway among different cell types. **(B)** The contribution of each L-R pair to the overall signaling pathway. **(C)** Violin plots showing the expression distribution of L-R gene pairs among different cell populations. **(D)** The network centrality analysis of *SPP1* pathway to investigate the out-degree, in-degree, flow betweenness and information centrality. **(E)** Circle plot. **(F)** L-R pairs contribution, **(G)** violin plots, and **(H)** network centrality analysis of *VISFATIN* pathway. **(I)** Circle plot, **(J)** L-R pairs contribution, **(K)** violin plots, and **(L)** network centrality analysis of *SEMA4* pathway.

**Figure S9.**
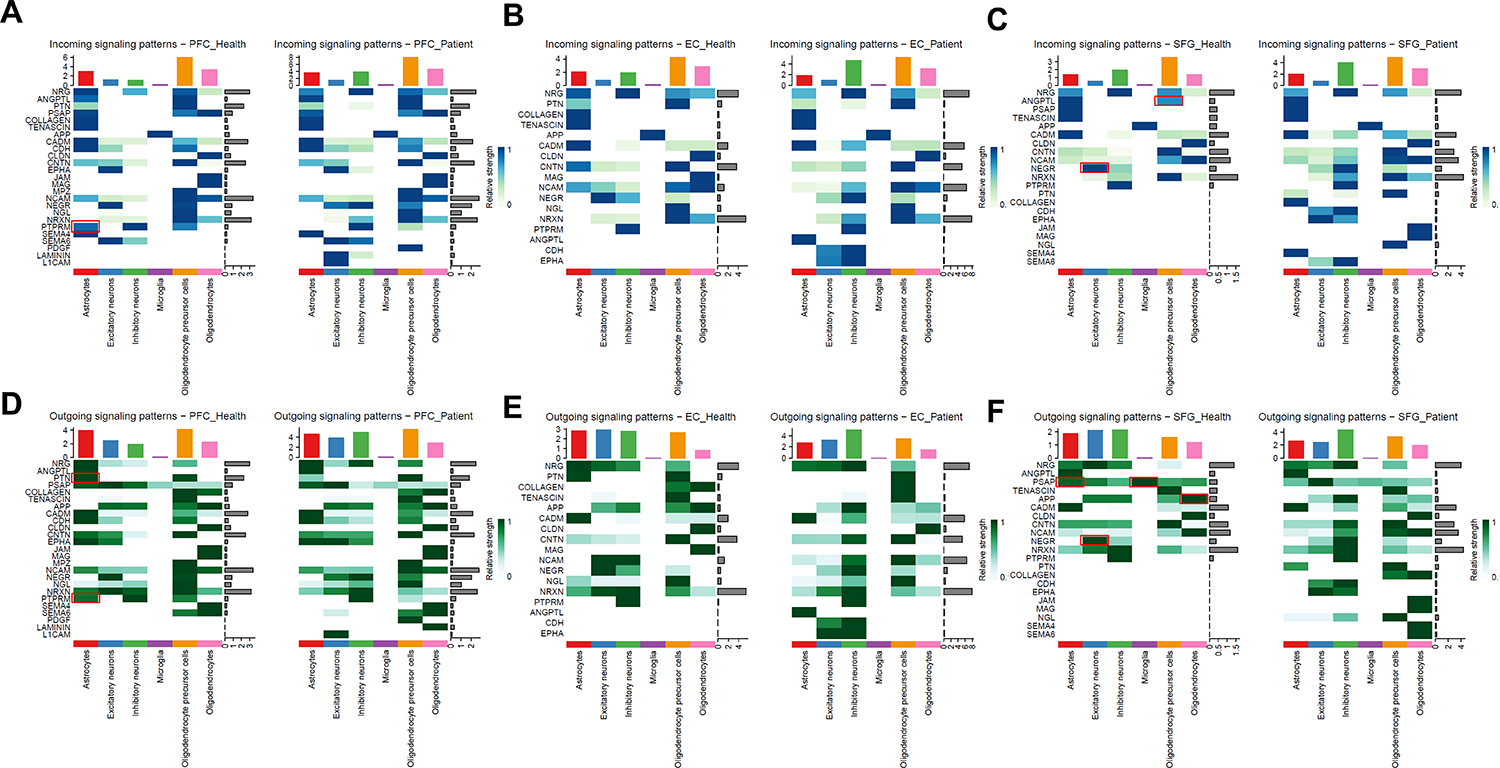
Summary of differential incoming and outgoing signaling patterns across six major cell types under different brain regions. **(A-C)** Incoming signaling patterns under **(A)** PFC, **(B)** EC, and **(C)** SFG regions. **(D-F)** Outgoing signaling patterns under **(D)** PFC, **(E)** EC, and **(F)** SFG regions.

**Figure S10.**
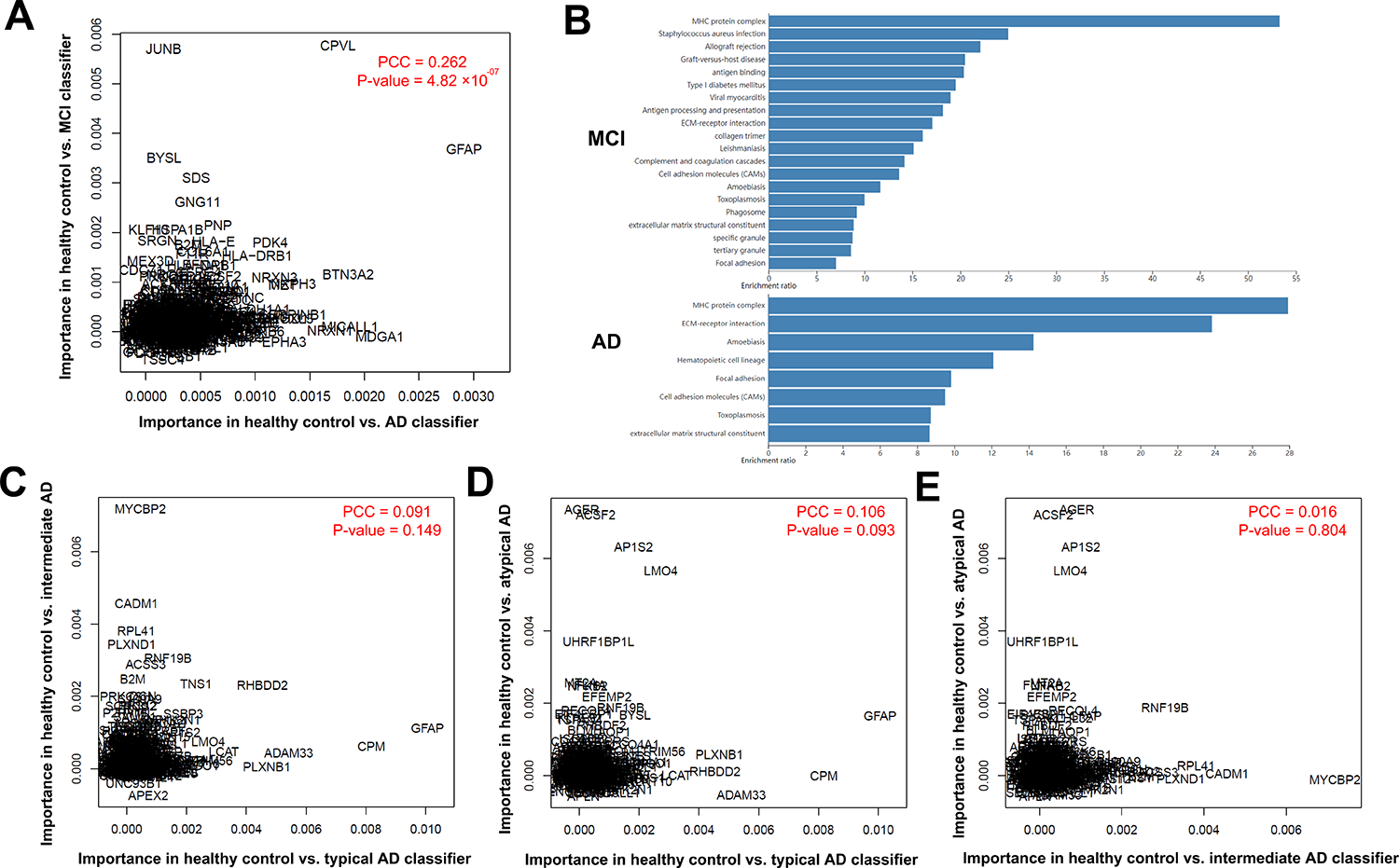
The feature comparison among different random forest models. **(A)** The feature importance comparison between model A (AD vs. healthy control) and model B (MCI vs. healthy control). The *X*-axis and *Y*-axis represent the mean decrease accuracy score in model A and model B, respectively. **(B)** GO and KEGG pathway enrichment analysis for high important genes (mean decrease accuracy > 0.005). **(C)** The feature importance comparison among model C (typical AD subtype vs. healthy control), **(D)** model D (intermediate subtype vs. healthy control), and **(E)** model E (atypical AD subtype vs. healthy control).

